# Contribution of the eye and of *opn4xa* function to circadian photoentrainment in the diurnal zebrafish

**DOI:** 10.1101/2022.07.30.500388

**Authors:** Clair Chaigne, Dora Sapède, Xavier Cousin, Laurent Sanchou, Patrick Blader, Elise Cau

## Abstract

The eye is instrumental for controlling circadian rhythms in mice and human. Here, we address the conservation of this function in the zebrafish, a diurnal vertebrate. Using lakritz (lak) mutant larvae, which lack retinal ganglion cells (RGCs), we show that while a functional eye contributes to masking, it is largely dispensable for the establishment of circadian rhythms of locomotor activity. Furthermore, the eye is dispensable for the induction of a phase delay following a pulse of white light at CT 16 but contributes to the induction of a phase advance upon a pulse of white light at CT21. Melanopsin photopigments are important mediators of photoentrainment, as shown in nocturnal mammals. One of the zebrafish melanopsin genes, *opn4xa*, is expressed in RGCs but also in photosensitive projection neurons in the pineal gland. Pineal *opn4xa*+ projection neurons function in a LIGHT ON manner in contrast to other projection neurons which function in a LIGHT OFF mode. We generated an *opn4xa* mutant in which the pineal LIGHT ON response is impaired. This mutation has no effect on masking and circadian rhythms of locomotor activity, or for the induction of phase shifts, but slightly modifies period length when larvae are subjected to constant light. Finally, analysis of *opn4xa;lak* double mutant larvae did not reveal redundancy between the function of the eye and *opn4xa* in the pineal for the control of phase shifts after light pulses. Our results support the idea that the eye is not the sole mediator of light influences on circadian rhythms of locomotor activity and highlight differences in the circadian system and photoentrainment of behaviour between different animal models.

**Author summary:** Experiments performed in mice have established a crucial role for the eye in general and melanopsin expressing cells in particular in the control of circadian rhythms most notably during photoentrainment, by which circadian rhythms adapt to a changing light environment. In marked contrast to this, we show that in zebrafish the eye and photosensitivity dependent on one of the melanopsin genes, *opn4xa*, which is expressed in both the eye and the pineal gland, are largely dispensable for correct circadian rhythms. These results provide insight that the light sensors orchestrating circadian rhythms of locomotor activity are different between animal models supporting that vertebrates might employ different molecular/cellular circuits for photoentrainment of behaviour depending on their phylogeny and/or temporal niche.

## INTRODUCTION

Light has a profound influence on the physiology and behaviour of living organisms. In particular, it controls circadian rhythms that in turn regulate a variety of biological functions. Circadian rhythms are defined by their period of approximately 24 hours. Once established, these rhythms persist in constant conditions, which has fostered the concept of an endogenous time-keeping mechanism known as the circadian system. Nonetheless, external cues are required to synchronize (or ‘entrain’) circadian rhythms with the exogenous environmental conditions. For instance, light entrains the circadian system through a process referred to as photoentrainment (see (Bhadra et al., 2017) for a review).

In mouse, photoentrainment depends on a functional retina. Eye enucleated mice or mice lacking retinal ganglion cells (RGCs) do not entrain to LD (Light/Dark) cycles and thus behave as if they were in constant darkness (Brzezinski et al., 2005; Freedman et al., 1999; Wee et al., 2002). Similarly, in human, fifty percent of blind people exhibit circadian misalignment with the LD cycles (Hartley et al., 2018; Lockley et al., 2007; Sack et al., 1992). In mouse, photoentrainment depends on a specific subtype of RGCs expressing the photopigment melanopsin, which is encoded by the Opn4 gene. These RGCs are sensitive to blue light and are referred to as ipRGCs for “intrinsically photosensitive RGCs”. Mice mutant for *Opn4* shows a diminished phase-delay in response to a pulse of light administered at circadian time 16 (CT 16; at the beginning of the subjective night) but entrain normally to LD cycles (Panda et al., 2002; Ruby et al., 2002). In contrast, mice with no ipRGCs or with impaired neurotransmission from ipRGCs show no entrainment to LD as well as no phase delay following a light pulse at CT16 (Gompf et al., 2015; Güler et al., 2008; Kofuji et al., 2016). The difference between the phenotypes observed when only melanopsin photosensitivity is impaired compared to the models where ipRGCs inputs to the brain are lost is thought to result from the influence of classical rods and cone photoreceptors on ipRGCs. Indeed, both rods and cones have been shown to play a role during photoentrainment and to signal to ipRGCs (Altimus et al., 2008; Belenky et al., 2003; Calligaro et al., 2019; Dkhissi-Benyahya et al., 2007; Dollet et al., 2010; Perez-Leon et al., 2006; Wong et al., 2007). Thus, ipRGCs function as a hub that integrates and transmits light information to the brain through a direct projection to the core of the suprachiasmatic nucleus (SCN; (Baver et al., 2008; Fernandez et al., 2016; Li and Schmidt, 2018)).This hypothalamic nucleus functions as a ‘master clock’ that synchronizes peripheral clocks present everywhere in the body.

In addition to photoentrainment, ipRGCs also control the increase of period length when animals are placed in constant light (LL) (Panda et al., 2002; Ruby et al., 2002) and are required for a process of maturation of the circadian clock that sets the definitive period of locomotor rhythms in LD and in constant darkness (DD, Chew et al., 2017). Finally, in addition to their crucial influence on the circadian system, murine ipRGCs also control masking, a direct suppressive effect of light on locomotor activity. This activity is thought to involve different ipRGC subtypes than the ones that impact the circadian system (Rupp et al., 2019).

ipRGCs are well established to mediate circadian and direct effects of light on behaviour in nocturnal mammals. Based on the conservation of the nervous system, observations made in human blind people and the description of ipRGC populations in the diurnal rodent Arvicanthis ansorgei as well as in human, ipRGCs are considered to play a similar role in diurnal mammals (Karnas et al., 2013; Mure, 2021). While the zebrafish has emerged as a powerful non-mammalian diurnal vertebrate model for chronobiology, the neural circuit controlling photoentrainment of behaviours has not been identified in this species. The observation that two species of cavefishes bearing eye degeneration : *Astyanax mexicanus* and *Phreatichthys andruzzii* do not show robust light-entrainable circadian locomotor activity rhythms could suggest that eye function is crucial for photoentrainment of locomotor rhythms in fishes (Cavallari et al., 2011). However, the observed circadian locomotor phenotypes could result from additional photoreceptive deficits. Indeed, *Phreatichthys andruzzii* present mutations in several opsin genes that are expressed in extra-ocular locations (Cavallari et al.. 2011). While the function of the eye regarding circadian rhythms of behaviours has not been addressed in fishes, in the zebrafish model, an additional level of complexity arises from the observation that all cell types are photosensitive (Carr and Whitmore, 2005; Vallone et al., 2004; Whitmore et al., 2000). This local photodetection could serve local functions (metabolism, transcription, cell cycle) or participate to the control of behaviors, although some level of central control is expected for orchestrating a complex process such as behavior. As such, the relative importance of central versus peripheral control for the photoentrainment of locomotor activity is still unclear.

To begin addressing how photoentrainment is controlled in zebrafish we first tested the role of the retina using *lakritz* (*lak*) mutants in which all RGCs fail to develop and as such no connection exist between the eye and the CNS (Kay et al., 2001). *lak* mutant larvae entrain to LD cycles and maintain rhythms of locomotor activity with a period similar to their control siblings in constant darkness (DD) but show subtle alterations in constant light (LL). While we detected no defect in phase shifting in response to a pulse of white light produced in the early subjective night (CT16) in *lak* -/- larvae, we observed a reduction of the phase shift induced upon a similar pulse of light at CT21 in *lak* mutants. The zebrafish possesses five melanopsin genes that are all expressed in the retina, including *opn4xa* and opn4b in larval RGCs (Kölsch et al., 2021; Matos-Cruz et al., 2011). In addition, melanopsin expression is detected in extra-retinal tissues. For instance, *opn4xa* is expressed in a subpopulation of projection neurons in the pineal gland. Interestingly these *opn4xa*+ projection neurons function in a LIGHT ON fashion while *opn4xa*-projection neurons function in a LIGHT OFF manner (Sapède et al., 2020). We engineered an *opn4xa* mutant in which the LIGHT ON response of the pineal is impaired; *opn4xa* -/- larvae successfully entrain to LD cycles and maintain rhythms of locomotor activity in constant conditions albeit with a reduction of period in LL. Pulses of white light at CT16 and CT21 induced similar phase shifts in *opn4xa* mutant and *opn4xa*/lak double mutant larvae compared to controls. Our results suggest that the function of the retina and the LIGHT ON response of the pineal gland are not absolutely required for circadian photoentrainment in zebrafish, thus further highlighting differences in the circadian system and circadian photoentrainment between mammals and zebrafish.

## RESULTS

### The zebrafish eye is dispensable for the establishment of circadian rhythms

Photoentrainment in mouse and human requires a functional eye. We took advantage of *lak* mutant larvae that lack RGCs to address whether this role for the eye is conserved in the diurnal zebrafish. Homozygous *lak* mutant larvae lack neuronal connections between the eye and the brain and do not display an optomotor response (Covello et al., 2020; Kay et al., 2001; Neuhauss et al., 1999). The gene mutated in *lak*, encoding for the bHLH transcription factor ATOH7 (ATH5) is expressed in the developing retina and nowhere else (Masai et al., 2000). The *lak* ^th241^allele we used bears a point mutation functions as a null allele which result in a total absence of both RGCs and a neural connection between the eye and the brain (Kay et al., 2001). We compared the locomotor activity of homozygous *lak* mutants with siblings (*lak*+/+ and *lak*+/-) in different illumination conditions. For each of these conditions, three independent experiments were performed. Within each independent experiment, the same number of homozygous mutants and control siblings were randomly selected and the mean of the three experiments was plotted. In this manner, the weight of each experiment within the final mean was identical between mutant and sibling populations.

In cycles of 14h light: 10h dark (hereafter referred to as LD), both siblings and *lak* homozygous mutant larvae exhibit rhythms of locomotor activity that are aligned with the LD cycles (Fig 1B). However, compared to sibling larvae, *lak* mutants show a specific reduction of activity during the day (Fig 1.B, supplemental Table 1). Sibling and *lak* mutant LD-entrained larvae placed in constant darkness (DD) demonstrate rhythms of locomotor activity with similar levels (Fig 1.C, supplemental Tables 2). In addition, the periods of the rhythms observed in DD did not significantly differ between the two populations (Fig 1D). The reduction in activity observed during the day in LD conditions thus does not affect the persistence of rhythmicity under free running conditions in DD.

**Figure 1:**
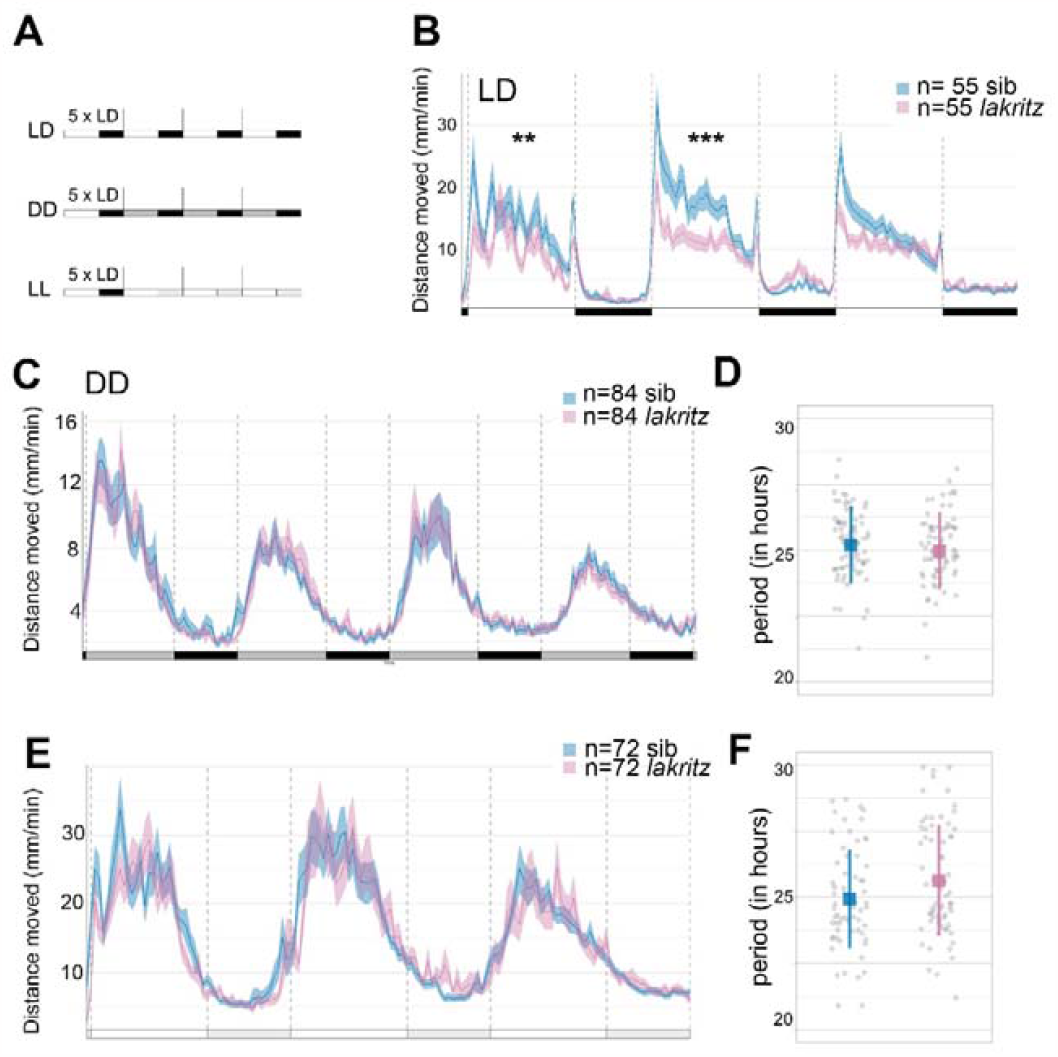
Locomotor activity of larvae devoid of RGCs in LD, DD and LL. **A)** Experimental design of LD, DD and LL experiments. White rectangles represent the day period, while black rectangles represent the night period, light grey rectangles represent the subjective day period and dark grey rectangles the subjective night. For each experiment, larvae are entrained for 5 LD cycles and their locomotor activity is tracked either in LD (LD), constant darkness (DD) or constant light (LL) the larvae are therefore 5dpf at the beginning of locomotor activity measurements. **B)** Average distance moved (mm/min) In LD. Merged data from 3 independent experiments represented in 10 min bins. Error bars represent SE. The distance moved is lower in *lak* larvae than control sibling larvae during the 1st (p=0.008) and 2nd days (p=0.005) but not during the 3rd day (p=0.13) nor during the night (p=0.42, p=0.51 and p=0.57 for the 1st, 2nd and 3rd nights; Mann-Whitney two-tailed test), see supplementary Table 1. * P<0.05, **p<0.01, ***p<0.005. **C)** Average distance moved in DD. Merged data from 3 independent experiments represented in 10 min bins. Error bars represent SE. No differences are detected between the distance moved of siblings versus *lak* larvae using a Mann-Whitney two-tailed test for each subjective night or day, see supplementary Table 2. **D)** Estimation of the periods using the FFT-NLLS method. Calculations were made on four complete cycles in DD. The mean period is not significantly different between sibling and *lak* larvae in DD (control: 25.08 ± 1.59 hours (n=114), *lak*: 24.95 ± 1.44 hours (n=83); mean ± S.D; p=0.32; Mann-Whitney two-tailed test, sibling vs *lak* larvae). Each grey point represents a single larva. **E)** Average distance moved in LL. Merged data from 3 independent experiments represented in 10 min bins. Error bars represent SE. **F)** Estimation of the periods using the FFT-NLLS method. Calculations were made on three complete cycles in LL. Mean± sd (in hours) is represented. Each grey point represents a larva, n= 66 siblings, n=68 *lak* larvae.

Finally, in constant light (LL) conditions, the activity of siblings and lak larvae were similar. The period of the lak mutant rhythm was slightly higher than the sibling rhythm but this difference was not significant (siblings: 24.93 ± 1.87 hours (n=66), lak: 25.63± 2.1 hours (n=68); mean ± S.D; p=0.083; Mann-Whitney). Altogether these results suggest that retinal ganglion cells and therefore a neuronal connection between the eye and the brain are dispensable for the establishment of circadian rhythms, their correct alignment to LD cycles and their maintenance in free running conditions (DD or LL).

### Absence of a functional eye differentially affects the induction of a phase delay and a phase-advance following pulses of light during the subjective night

To evaluate the role for RGCs in circadian photoentrainment, we assessed the phase-shifting effect of a pulse of white light on locomotor activity in *lak* larvae during the subjective night. We first chose to perform such a light pulse at CT16, as this was previously shown to induce a robust phase shift of the molecular clock in cell cultures (Tamai et al., 2007; Vallone et al., 2004). After entraining for 5 LD cycles, larvae from *lak*+/-incrosses were further reared in DD and subjected to a pulse of light during the second night in DD (“PD larvae”). Their activity was compared with the activity of larvae kept in the dark for 4 days (“DD larvae”). To analyze if a phase shift was induced, we calculated the difference of phase between the two last days (“after the light pulse”) and the two first days (“before the light pulse”), a value we refer to as “Δphase” (Fig 2.A). We found that a 2-hours pulse of light at CT16 induced a phase delay of locomotor activity rhythms in larvae, as the Δphase of PD larvae was higher than the one of DD larvae (Fig 2.B, Table 1). When the difference between the Δphase of pulsed larvae minus the Δphase of larvae placed in DD was calculated, it suggests a phase delay of 2.94 hours on average in PD larvae. Finally, we determined that control and *lak* mutant larvae exhibit a similar phase shift in locomotor activity (Fig 2.C, Table 1) suggesting that RGCs are not necessary for the circadian photoentrainment of locomotor activity to a pulse of light at the beginning of the subjective night (CT16).

**Figure 2:**
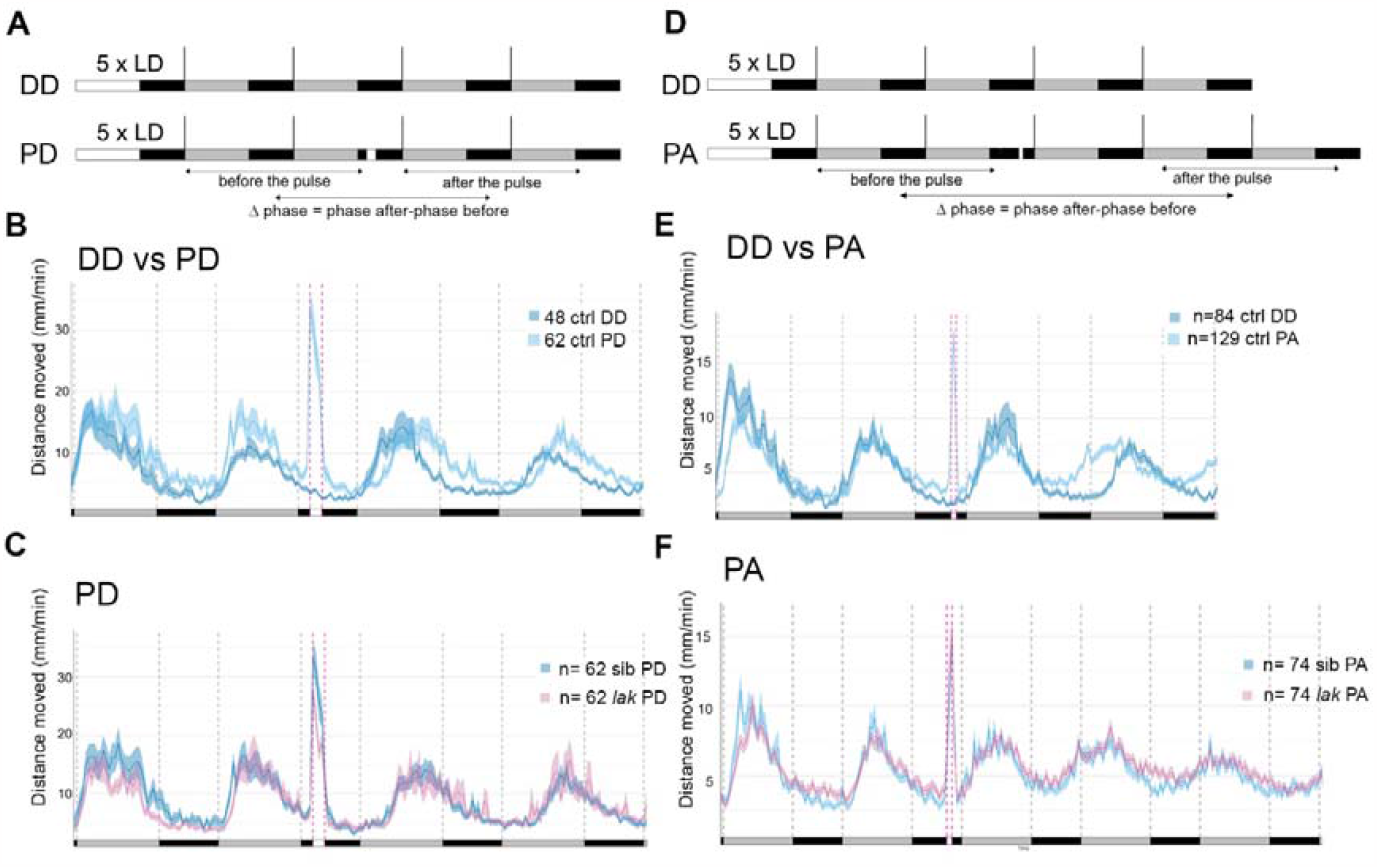
Larvae devoid of RGCs still photoentrain to pulses of white light at CT16 and CT21. **A)** Experimental design of phase delay (PD) experiments. White rectangles represent the day or light pulse period, black rectangles represent the night period and dark grey rectangles represent the subjective day. For each experiment, larvae are entrained for 5 LD cycles, the larvae are therefore 5dpf at the beginning of locomotor activity measurements. Locomotor activity is tracked either in constant darkness for 4 days (DD) or tracked in constant darkness for 4 days and subjected to a 2-hours pulse of light during the night of the 2nd day of constant darkness at CT16 (PD). The phase of locomotor activity is calculated for each larva before and after the timing of the pulse for DD and PD experiments and the Δphase (phase after the pulse – phase before the pulse) is calculated. **B)** Average distance moved by control larvae (mm/min over 10min) in DD and PD experiments. Mean ± SE. The Δphase calculated using the FFT-NLLS method of PD larvae is higher than the one of DD larvae (p<0.0001, Mann-Whitney two-tailed test), showing that the pulse of light induced a phase delay. **C)** Average distance moved merged from PD experiments represented in 10 min bins. Mean ± SE. The Δphase of control versus *lak* larvae calculated using the FFT-NLLS method is not significantly different (p=0.24, Mann-Whitney two-tailed test). *lak* show lower levels of activity during the light pulse (p=0.03, Mann-Whitney two-tailed test). **D)** Experimental design of phase advance (PA) experiments. The iconography is similar to A). PA-pulsed larvae were subjected to a one-hour pulse of light at CT21. **E)** Average distance moved of control larvae (mm/min over 10min) in DD and PD experiments Mean ± SE. The Δphase calculated using the FFT-NLLS method is negative in PA-pulsed larvae and statistically different from DD larvae (p<0.0001, Mann-Whitney two-tailed test), showing that the pulse of light induced a phase advance. **F)** Average distance moved merged from PA experiments represented in 10 min bins. Mean ± SE.

**Table 1:**
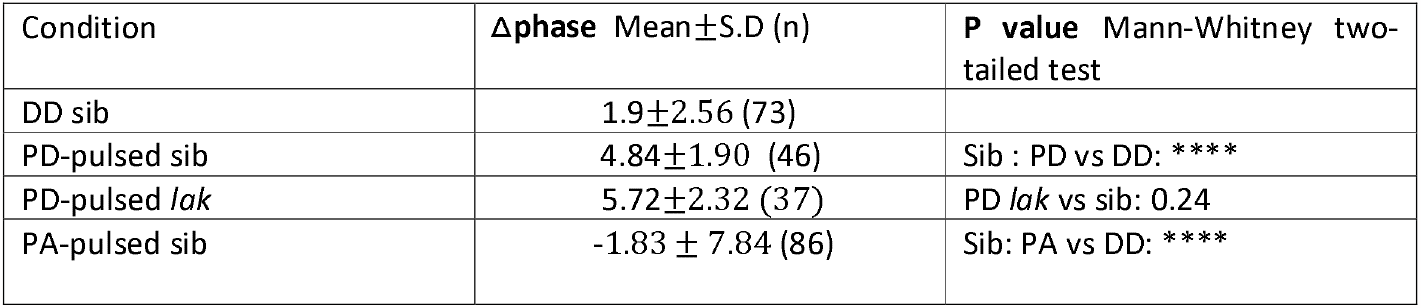
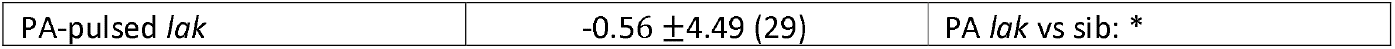
Quantification of the phase shifts in control siblings versus *lak*-/- (*lak)* larvae kept in DD or submitted to pulses of white light at CT16 or CT21.

| Condition | Δphase Mean ± S.D (n) | P value Mann-Whitney two-tailed test |
| --- | --- | --- |
| DD sib | $1.9 \pm 2.56$ (73) | |
| PD-pulsed sib | $4.84 \pm 1.90$ (46) | Sib : PD vs DD: **** |
| PD-pulsed *lak* | $5.72 \pm 2.32$ (37) | PD *lak* vs sib: 0.24 |
| PA-pulsed sib | $-1.83 \pm 7.84$ (86) | Sib: PA vs DD: **** |
| PA-pulsed *lak* | -0.56 ±4.49 (29) | PA *lak* vs sib: * |

**Table 2:** Quantification of the phase shifts in *opn4xa*+/+ versus *opn4xa*-/- larvae kept in DD or submitted to a 2 hours pulse of white light at CT16.

| Condition | Δphase Mean±S.D (n) | P value Mann-Whitney two-tailed test |
| --- | --- | --- |
| DD *opn4xa* +/+ | 1.64±2.92 (39) | |
| PD-pulsed *opn4xa* +/+ | 4.73±2.63 (36) | *opn4xa* +/+ : PD vs DD: **** |
| PD-pulsed *opn4xa* -/- | 5.23±4.81 (35) | PD *opn4xa* -/- vs *opn4xa* +/+ : 0.32 |
| PA-pulsed *opn4xa* +/+ | -2.61±4.05 (60) | *opn4xa* +/+ : PA vs DD: **** |
| PA-pulsed *opn4xa* -/- | -2±2.88 (20) | PA *opn4xa* -/- vs *opn4xa* +/+ : 0.39 |

We next performed a pulse of light at the end of the subjective night. A one hour pulse induced a phase advance that could be detected during the second circadian cycle after the pulse (Figure 2D, E). Since this phase shift is not seen during the first cycle, we analyzed larvae over an additional cycle in order to obtain enough data to perform a robust phase calculation. The difference between the Δphase of pulsed larvae (PA) minus the Δphase of larvae placed in DD alone suggests that this paradigm induced a phase advance of at least 3.7 (1.8 +1.9) hours on average. While *lak* -/- larvae showed a phase advance upon a pulse of light at CT21 (Fig 2.F, table1), this phase shift was weaker than that induced in control larvae (Δphase= -1.8 ± 7.84 for siblings versus Δphase = -0.56 ± 4.49 for *lak* -/- larvae). Although this difference in phase shift between *lak* and siblings is difficult to observe on the graph representing all the animals from the four independent experiments, it better shows when only the animals for which a phase was successfully extracted before and after the pulse (Figure 2F, Figure S1). These results suggest that a phase advance can occur in absence of RGCs although the eye contributes to photoentrainment in this context. Altogether, our results thus suggest that although phase advances and delays can occur in absence of RGCs, the absence of these cells specifically affect the response to a phase advance paradigm.

The Δphase is the difference between the phase of the two last cycles and the phase of the two first cycles. A phase shift is observed in DD owing to the period that is close to 25 hours which generates a ~1 hour-shift every cycle. Upon a pulse of light at CT16 or CT21 a statistical difference is observed between DD and pulsed sib larvae as well as *lak* and sib larvae when the pulse of light is applied at CT21 (****, p<0,0001; *, p<0,05 using a Mann-Whitney two-tailed test). For each type of paradigm, (DD, PD and PA) three independent experiments were pooled.

To analyse a role for RGCs during masking we calculated the activity of control and lak larvae during the pulses of light performed at CT16 and CT21. Interestingly, *lak* larvae showed a reduced level of activity compared to control larvae during the pulse performed at CT16 but not at CT21 (CT16 : Fig 2.C, siblings: 28.33 ± 19.26 mm/min over 10min (n=62), *lak*: 21.45 ± 12.27 mm/min over 10min (n=51); p=0.03; Mann-Whitney two-tailed test; CT21 : Fig 2.F siblings: 16.51 ± 10.9 mm/min over 10min (n=51), *lak*: 17.17 ± 7.85 mm/min over 10min (n=62); p=0.75). These results showed that RGCs are involved in masking in the zebrafish larvae but in a circadian dependent manner.

### opn4xa function contributes to endogenous period setting in LL

As circadian rhythms of locomotor activity are established and photoentrain in absence of RGCs, albeit with less efficiency, we wondered if *opn4xa*+ projection neurons present in the pineal gland could play a role in the establishment and photoentrainment of circadian rhythms (Sapède et al., 2020). We, thus, generated a mutant allele for *opn4xa* via CRISPR/Cas9 genome editing using guide RNAs targeting the second coding exon. Amongst various alleles that were generated, we selected an allele that displays a 17 nucleotides insertion for further analysis (Fig 3.A). The protein encoded from this allele is predicted to contain a premature stop codon (Fig S2.A) leading to a truncation of the protein in the middle of the second transmembrane domain (Fig S2.B), which should result in a protein devoid of a G protein interaction domain. We have verified that the two alternative ORFs also generate a truncation from the mutated allele (Fig S2.C) and that the use of an alternative ATG present in both the wt and the mutant allele also leads to a truncation (Fig S2. D). Finally, we have used Spliceator as to verify that the mutation does not generate alternative splicing sites (Spliceator (lbgi.fr)). We therefore predicted production of a null allele. Homozygous animals were viable and fertile.

**Figure 3:**
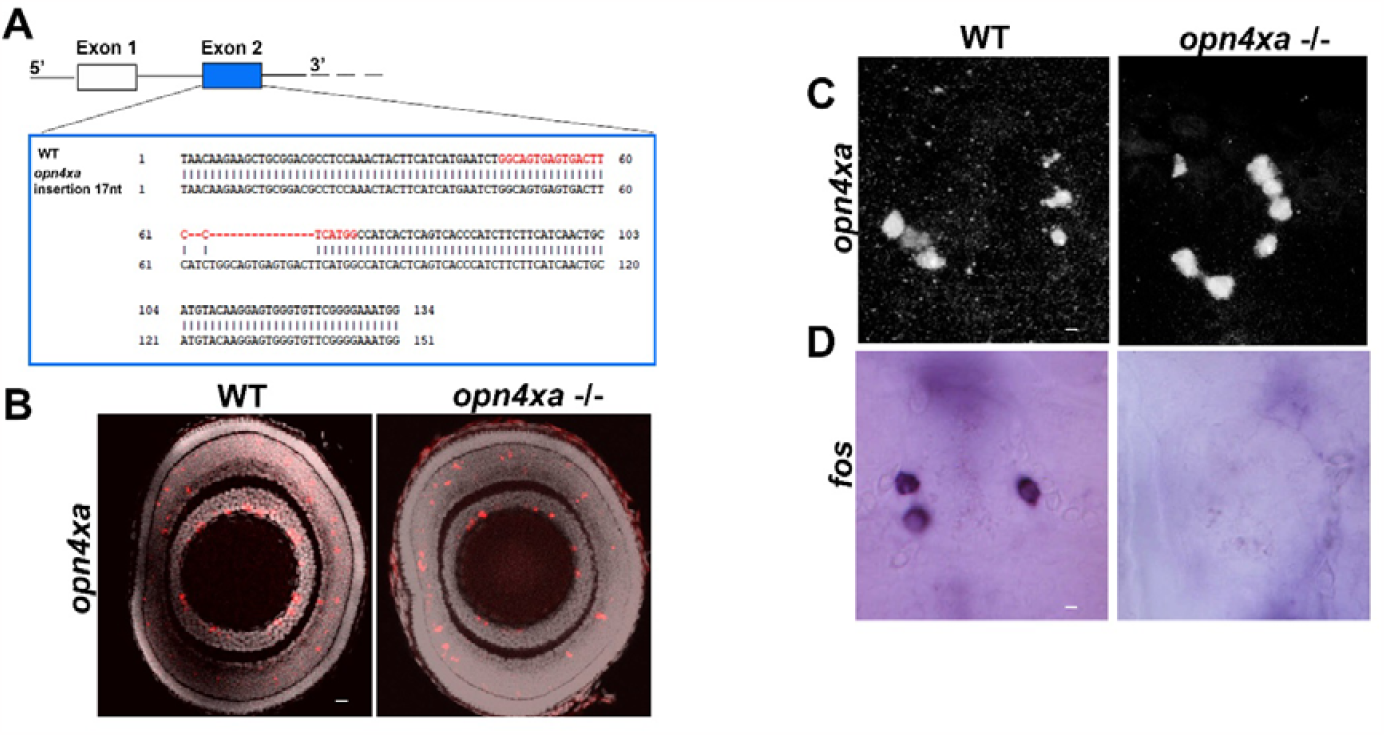
Mutation in *opn4xa* abolishes light sensitivity in pineal *opn4xa*+ cells. (A) Scheme showing the 5’ part of the *opn4xa* locus and in particular the second exon targeted by the CRISPR guide RNA (target sequence is highlighted in red) as well as the WT and mutant exon2 sequences. B) In situ hybridization showing *opn4xa* expression in the retina at 4 days (ZT0) (C) Expression of *opn4xa* in the pineal at 4 days (ZT0) (D) fos expression at 3 days after 30 min of illumination. Scale bars respectively represent 10 µm (B) and 5 µm (C-D).

We first looked at the expression of *opn4xa* in the retina of *opn4xa*+/+ and *opn4xa* -/- larvae. The RGC layer contained an average of 33 to 44 *opn4xa*+ cells (Fig S3 A-E), which number did not significantly vary between the different time points suggesting an absence of diel rhythm in this layer. Surprisingly, we observed a previously undescribed expression of *opn4xa* in the interneuron layer with numerous cells at 4dZT0 and 5dZT0 and very few cells at the other ZT time points (Fig S3A-D, F). The number of *opn4xa*+ cells in both the RGC and the interneuron layer are normal in the *opn4xa* -/- retina (Fig 3B, RGC layer : *opn4xa* +/+: 26.17 ± 7.33, n= 6; *opn4xa*-/-: 23.8±6.92, n=10; p=0.42 using a Mann-Whitney two-tailed test; interneurons layer : *opn4xa* +/+: 26.33± 11.5 ; *opn4xa*-/-: 22.9±7.19; p=0.63 using a Mann-Whitney two-tailed test). Similarly, we have observed that the number of *opn4xa* + pineal cells were similar in *opn4xa* +/+ and *opn4xa* -/- larvae (Fig 3 C, *opn4xa* +/+: 4.33±1.51 n=6, *opn4xa*-/- : 4.75± 1.98 n=9, p=0.85 using a Mann-Whitney two-tailed test). In addition, pineal *opn4xa*+ cells express the Wnt effector tcf7 (Sapède et al., 2020). At 6 days post fertilization, the *opn4xa*-/- pineal gland displayed normal expression of tcf7 (opn4xa +/+: 5.5 ± 2,5 (n=4), *opn4xa*-/-: 6± 3 cells (n=3); mean ± S.D; Figure S4). Upon illumination with a 30 min pulse of light, fos is expressed in 2-5 cells of the pineal gland (Fig 3.D). We have previously shown that this expression corresponds to *opn4xa*+ cells (Sapede et al., 2020). On the other hand, fos expression is virtually absent in the pineal gland of *opn4xa*-/- embryos after a 30 min pulse of light (Fig 3.D; at 3 days *opn4xa* +/+: 4.8 ± 0.8 (n=9), *opn4xa*-/-: 0.25 ± 0,7 cells (n=8) per pineal; at 7 days *opn4xa* +/+: 5.9 ± 3,5 (n=9) *opn4xa*-/-: 0.4 ± 0,7 cells (n=13); mean ± S.D). Altogether our results suggest that *opn4xa*-/- larvae contained normal numbers of *opn4xa*+ cells in the retina and the pineal gland but have lost expression of fos following a 30 mn light pulse. We therefore used the *opn4xa* mutant as a model in which the pineal ON_response is impaired.

To identify a role for *opn4xa* in the control of circadian rhythms, we analysed locomotor behaviour of *opn4xa*-/- larvae under various illumination regimes. Similar to the analysis we performed in lak larvae, for each of these conditions, three independent experiments were performed. Within each independent experiment, the same number of *opn4xa* -/- and *opn4xa* +/+ siblings were randomly selected and the mean of the three experiments was plotted. We found that *opn4xa* -/- larvae still entrained to LD cycles and did not show any difference in levels of locomotor activity when compared to their *opn4xa* +/+ siblings (Fig 4.B + supplemental Table 4). In addition, *opn4xa*-/- larvae were able to maintain rhythms of locomotor activity with a similar period as wild-type larvae in DD (Fig 4.C, D). *opn4xa*-/- larvae placed in LL still showed circadian rhythms of locomotor activity (Fig 4.E, F) but with several alterations. First, *opn4xa*-/- larvae were significantly more active during the first night (Supplemental table 6). More importantly, in LL the period was shorter for *opn4xa*-/- larvae compared to controls using both the FFT-NLLS (opn4xa+/+: 25.31 ± 3.29 hours (n=65), *opn4xa*-/-: 24.71 ± 3.32 hours (n=64); p=0.041; Mann-Whitney two-tailed test) and mFourfit methods (opn4xa+/+: 25.84 ± 1.60 hours (n=66), *opn4xa*-/-: 25.12 ± 1.739 hours (n=66); p=0.012). Altogether these results suggest that *opn4xa* contributes to endogenous period setting in LL.

**Figure 4:**
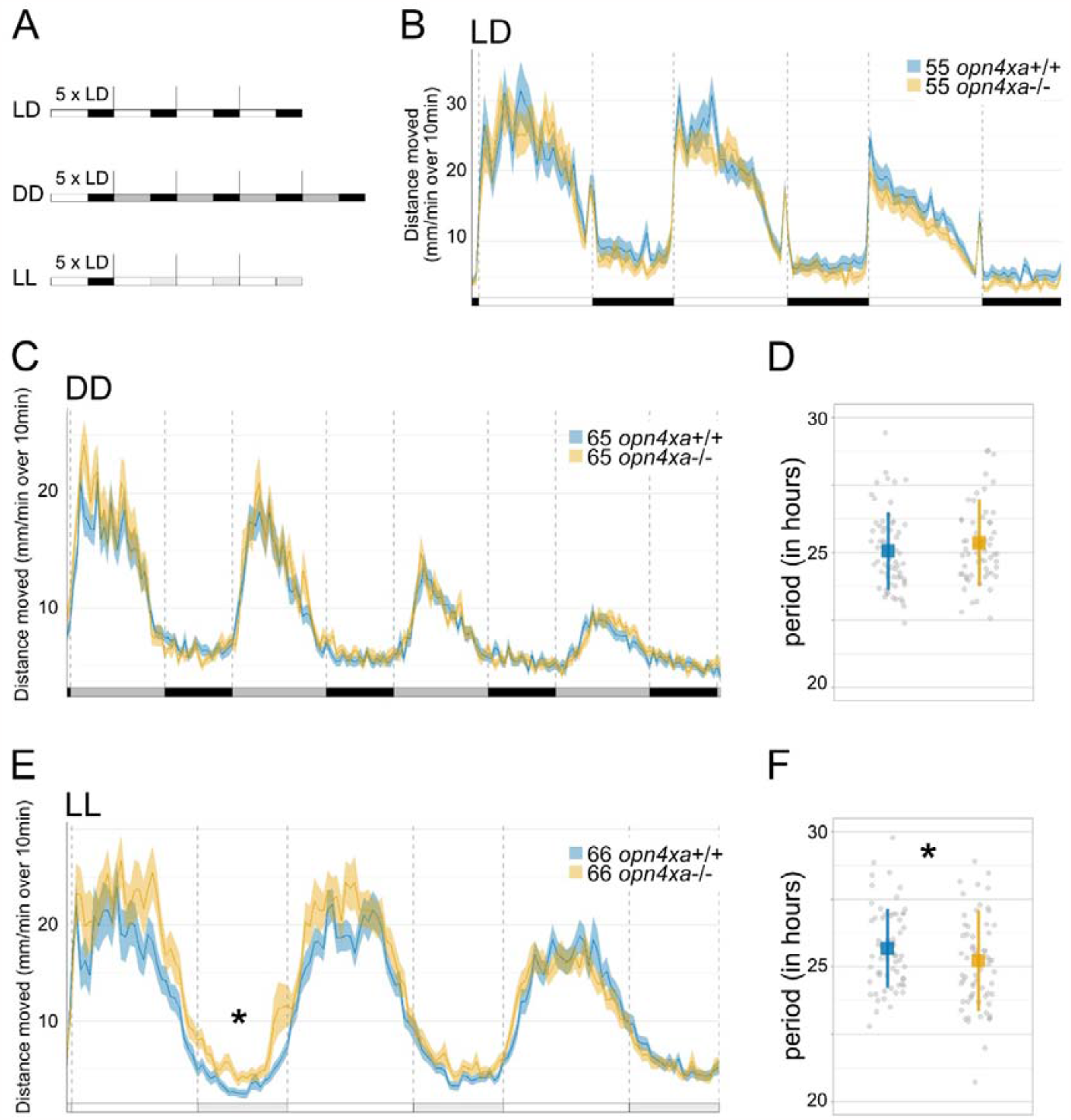
Locomotor activity of larvae devoid of *opn4xa*-mediated photosensitivity (opn4xa-/-) in LD, DD and LL. **A)** Experimental design of LD, DD and LL experiments. White rectangles represent the day period, black rectangles represent the night period, dark grey rectangles represent the subjective day period and light grey rectangles the subjective night. For each experiment, larvae are entrained for 5 LD cycles and their locomotor activity is tracked either in LD (LD), constant darkness (DD) or constant light (LL) the larvae are therefore 5dpf at the beginning of locomotor activity measurements. **B)** Average distance moved (mm/min over 10min) of 3 independent experiments in LD. Mean ± SE. The distance moved is not different in *opn4xa*+/+ and *opn4xa*-/-larvae during the 1st (p=0.73), 2nd days (p=0.50) and 3rd days (p=0.07) nor during the 1st (p=0.30), and 2nd nights (p=0.27) (supplemental Table 4). A lower level of activity is found in *opn4xa*-/- larvae during the 3rd night (p=0.01) but is visually clear in only one of the 3 independent experiments (Mann-Whitney two-tailed test). **C)** Average distance moved (mm/min over 10min) of 3 independent experiments in DD. Mean ± SE. **D)** Estimation of the periods using the FFT-NLLS method calculated over four cycles. The mean period is not significantly different between control and *opn4xa*+/+ and *opn4xa*-/- larvae in DD (opn4xa+/+: 25.05 ± 1.43 hours (n=64), *opn4xa*-/-: 25.35 ± 1.60 hours (n=60); mean±SD; p=0.29; Mann-Whitney two-tailed test). Mean± sd (in hours) is represented. Each grey point represents a larva. **E)** Average distance moved (mm/min over 10min) of 4 independent experiments in LL. Mean ± SE. *opn4xa*-/- are more active than controls during the first night (p=0.02, see supplemental Table 6). **F)** Estimation of the periods using the FFT-NLLS method calculated over three cycles. The mean period is significantly different between *opn4xa*+/+ and *opn4xa*-/- larvae in LL. Mean± sd (in hours) is represented. Each grey point represents a larva.

Since *opn4xa*-/- larvae showed a slight hyperactivity during the first night as well as a subtle period phenotype in LL, we began to analyse the possible molecular mechanisms behind these effects using RTqPCR (Figure 5). Analysis of clock gene expression in *opn4xa*-/- versus wt background reveal normal rhythms of clock genes expression in LD except for a slight increase in tef alpha and cry1a in *opn4xa* -/- larvae (Figure 5 B,D). In LL after training, the expression of bmal1a was higher in *opn4xa* -/- than in wt larvae (Figure 5A). Interestingly, the expression of some Bmal1a direct targets (cry1a, dec1) was also increased (figure 5 B, C) while expression of reverbB1, per1a, per1b, per2 remain unchanged (Figure 5 E-H). Overall, only subtle and specific molecular phenotypes were detected in *opn4xa* -/- larvae in LL. In addition, their relation to the locomotor activity phenotypes are not clear.

**Figure 5:**
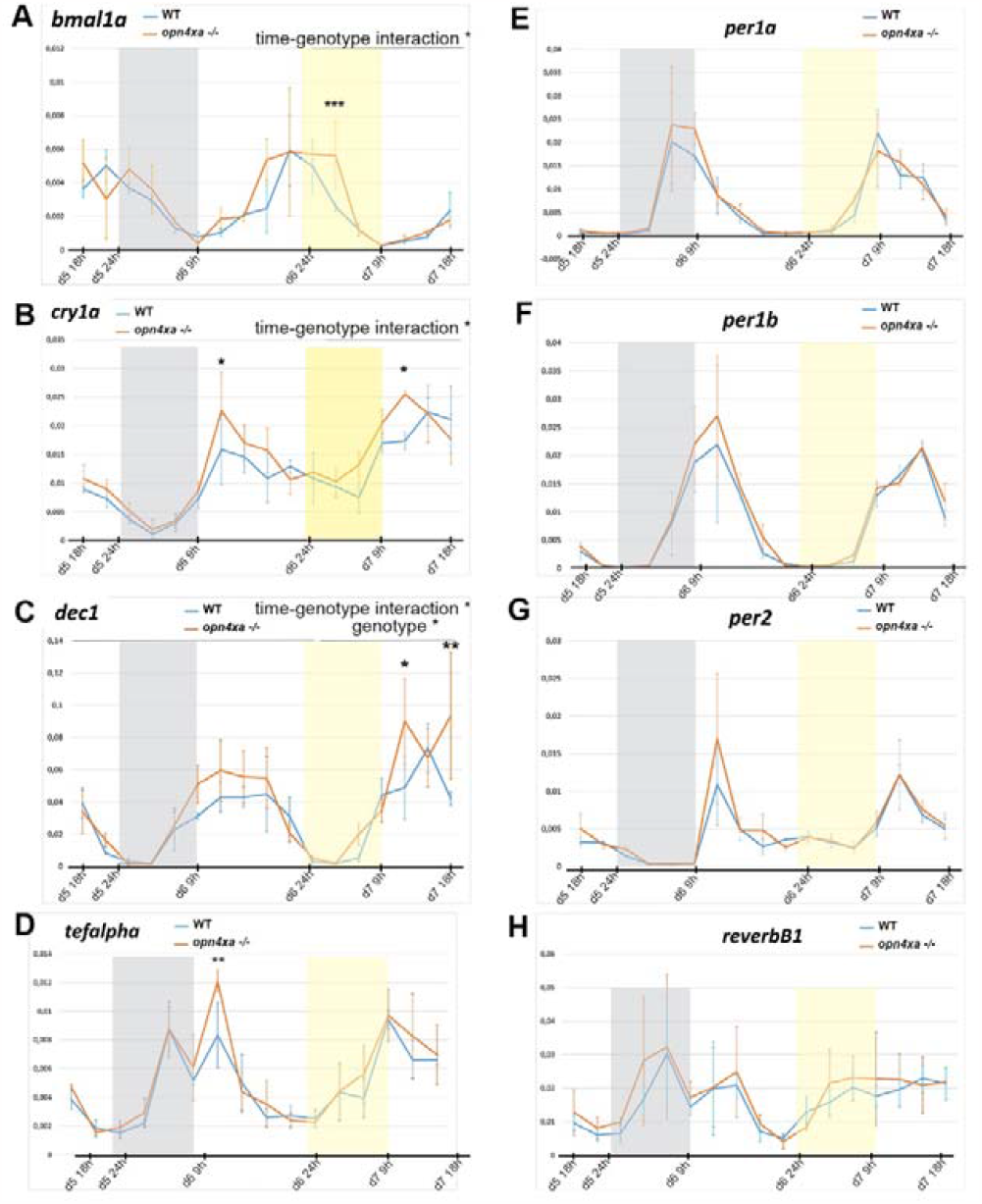
opn4xa -/- larvae show subtle modifications of a few clock genes in LL : RTqPCR performed on pools of 15 larvae for the gene indicated at the top of the figures. Mean expression relative to beta actin ± s.d. Three pools of larvae were used for each time point. ‘wt’ refers to pool of larvae from crosses of *opn4xa*+/+ animals (siblings of the *opn4xa*-/- fishes used for the *opn4xa*-/-points). Larvae were exposed to LD cycles (until d6 21h) followed by a LL cycle (from d6 21h to d7 18h). The grey rectangles represent the night phase, the yellow rectangles represent the subjective night in the LL cycle. The data were analysed using two way ANOVAs which revealed time-genotype interaction for *bmal1a* and *cry1a*, as well as statistical differences between genotypes for specific time points using Bonferroni post-hoc tests. p< 0.05; ** p< 0.001; ***p< 0.0005.

### opn4xa function is dispensable for photoentrainment to a pulse of white light during the subjective night

We next assessed the ability of pulses of white light at CT 16 and CT21 to induce phase shifts in an *opn4xa*-/- background. We observed that under such conditions *opn4xa*-/- larvae shift their activity to the same extent as their wild-type siblings (Fig 6, Table 3). Furthermore, *opn4xa*-/- larvae did not show any difference in the level of activity during the pulses of light at CT16 or CT21 compared to wildtype siblings, implying that photosensitivity controlled by *opn4xa* is not required for masking (CT16, Fig 6.B, *opn4xa*+/+: 20.30 ± 10.03 mm/min over 10min (n=58), *opn4xa*-/-: 19.42 ± 10.76 mm/min over 10min (n=58); p=0.56; Mann-Whitney two-tailed test. CT21: Fig 6.D, *opn4xa*+/+: 12.85 ± 6.80 mm/min over 10min (n=44), *opn4xa*-/-: 16.54 ± 8.55 mm/min over 10min (n=44); p=0.47). These results show that the intrinsic photosensitivity of *opn4xa* expressing cells is not necessary for circadian photoentrainment or masking.

**Table 3:** Quantification of the phase shifts in *lak* versus double mutant larvae kept in DD or submitted to pulses of white light at CT16 or CT21.

| Condition | $\Delta\text{phase}$ Mean ± S.D (n) | P value Mann-Whitney two-tailed test |
| --- | --- | --- |
| DD *lak* | 2.143 ± 2.09 (31) | |
| PD-pulsed *lak* | 5.72 ± 2.32 (37) | *lak*: PD vs DD: **** |
| PD-pulsed double | 4.90 ± 2.23 (14) | PD double vs *lak*: 0.33 |
| PA-pulsed *lak* | -0.57 ± 4.49 (29) | *lak*: PA vs DD: **** |
| PA-pulsed double | -3.16 ± 4.9 (16) | PA double vs *lak*: 0.24 |

**Figure 6:**
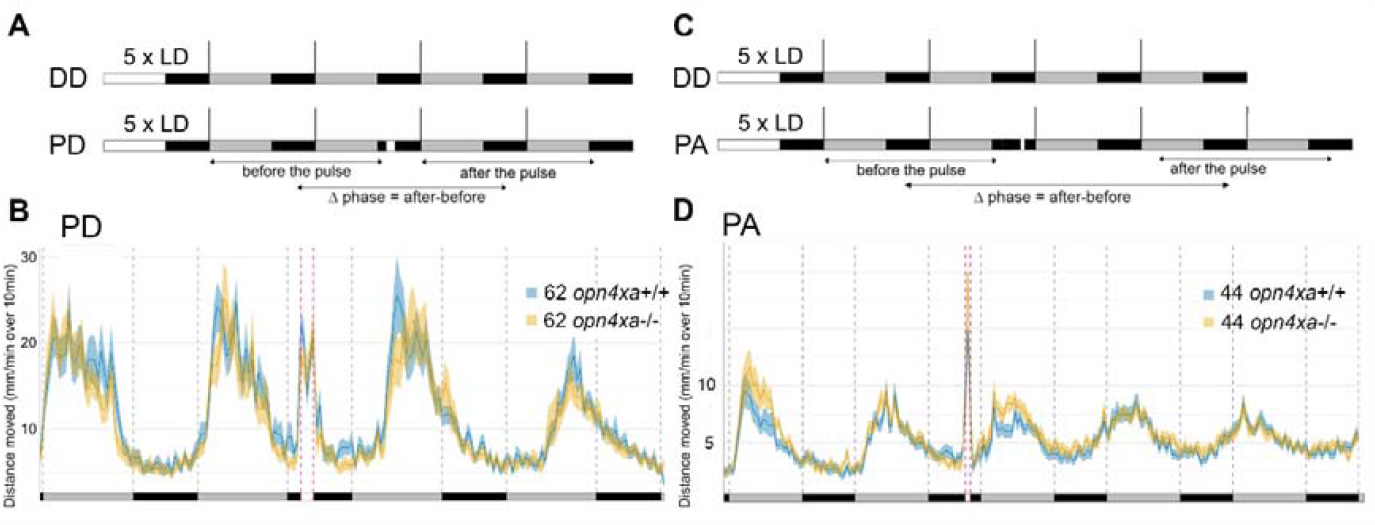
Larvae devoid of *opn4xa*-mediated photosensitivity (*opn4xa-/-*) still photoentrain to pulses of light at CT16 and CT21. **A)** Experimental design of phase shift experiments. White rectangles represent the day or light pulse period, black rectangles represent the night period and dark grey rectangles represent the subjective day. For each experiment, larvae are entrained for 5 LD cycles the larvae are therefore 5dpf at the beginning of locomotor activity measurements. Locomotor activity is tracked either in constant darkness for 4 days (DD) or tracked in constant darkness for 4 days and subjected to a 2-hours pulse of light during the night of the 2nd day of constant darkness (PD). The phase of locomotor activity is calculated for each larva before and after the timing of the pulse for DD and PS experiments and the Δphase (phase after the pulse – phase before the pulse) is calculated. **B)** Average distance moved (mm/min over 10min) of 3 independent PD experiments. Mean ± SE. The Δphase of *opn4xa*+/+ and *opn4xa*-/- larvae calculated with the FFT-NLLS method is not significantly different. *opn4xa*+/+ and *opn4xa*-/-show similar levels of activity during the light pulse (p=0.56, Mann-Whitney two-tailed test). **C)** Experimental design of phase advance (PA) experiments. The iconography is similar to A). PA-pulsed larvae were subjected to a one-hour pulse of light at CT21. **D)** Average distance moved (mm/min over 10min) of 3 independent PA experiments. Mean ± SE. The Δphase of *opn4xa*+/+ and *opn4xa*-/- larvae calculated with the FFT-NLLS method is not significantly different (Mann-Whitney two-tailed test).

As for table 1, the Δphase is the difference between the phase of the two last cycles and the phase of the two first cycles. A Phase shift was observed in DD owing to the period that is close to 25 hours which generates a ~1 hour-shift every cycle. Phases were calculated with the FFT-NLLS method. Upon a pulse of light at CT16 or CT21 a statistical difference was observed between DD and pulsed ctrl larvae (****, p<0.0001 using a Mann-Whitney two-tailed test). In both phase delays (PD, pulse of light at CT16) and phase advance paradigms (PA, pulse of light at CT21), no statistical difference between *opn4xa*+/+ and *opn4xa*-/- was observed using a Mann-Whitney two-tailed test. For each type of paradigm (DD, PD and PA) three independent experiments were pooled.

Since neither the absence of RGCs (Fig 2) nor the loss of *opn4xa*-dependent photosensitivity (Fig 6) abolished the capacity of larvae to photoentrain to pulses of light performed in the early or late subjective night, a possible compensation could occur between RGCs and *opn4xa* expressing cells of the pineal gland. To begin addressing this question, we tested photoentrainment properties of lak-/-; *opn4xa*-/- larvae (referred to as ‘double’). Compared to lak simple mutants, double mutants did not show an attenuated phase shift response to pulses of light at CT16 or CT21 (Fig 7). This suggests that other photosensitive cells mediate photoentrainment in zebrafish.

**Figure 7:**
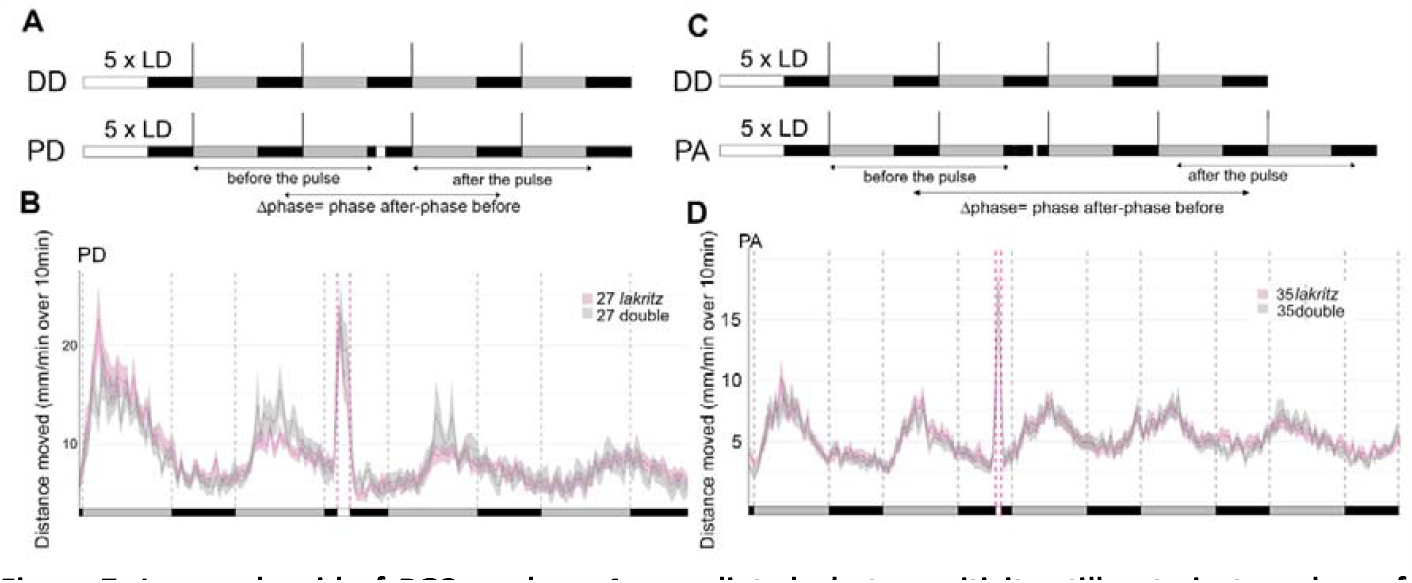
Larvae devoid of RGCs and *opn4xa*-mediated photosensitivity still entrain to pulses of light at CT16 and CT21. **A)** Experimental design of phase shift experiments. White rectangles represent the day or light pulse period, black rectangles represent the night period and dark grey rectangles represent the subjective day. For each experiment, larvae are entrained for 5 LD cycles the larvae are therefore 5dpf at the beginning of locomotor activity measurements. Locomotor activity is tracked either in constant darkness for 4 days (DD) or tracked in constant darkness for 4 days and subjected to a 2-hours pulse of light during the night of the 2nd day of constant darkness (PD). The phase of locomotor activity is calculated for each larva before and after the timing of the pulse for DD and PS experiments and the Δphase (phase after the pulse – phase before the pulse) is calculated. **B)** Average distance moved (mm/min over 10min) of 3 independent PD experiments (n=27 for *lak* -/- referred as *lak* and n=27 *lak*-/-; *opn4xa*-/- larvae referred as ‘double’). Mean ± SE. The Δphase of *lak* and double larvae calculated with the FFT-NLLS method are not significantly different (see table 3). **C)** Experimental design of phase advance (PA) experiments. The iconography is similar to A). After 5 training cycles in LD, PA-pulsed larvae were subjected to a one-hour pulse of light at CT21. **D)**Average distance moved (mm/min over 10min) of 3 independent PA experiments (n=27 for *lak* and n=27 double larvae). Error bars represent SE. The Δphase of *lak* and double larvae calculated with the FFT-NLLS method are not significantly different (see table 3).

Finally, *lak* and *lak*/opn4xa double mutant larvae show similar levels of activity during the light pulse both at CT 16 and CT21 (CT16: lak : 18.23 ± 8.65 mm/min over 10min (n=27), double: 19.79 ± 12.39 mm/min over 10min (n=27); p=0.94, CT21 : *lak*: 17.46 ± 6.65 mm/min over 10min (n=27), double: 16.8 ± 8.05 mm/min over 10min (n=27), p=0.42, Mann-Whitney two-tailed test). This suggests an absence of redundancy between RGC and *opn4xa* photosensitivity for masking of locomotor activity.

As for table 1 and 2, the Δphase is the difference between the phase of the two last cycles and the phase of the two first cycles. A Phase shift is observed in DD owing to the period that is close to 25 hours which generates a ~1 hour-shift every cycle. Phases were calculated with the FFT-NLLS method (biodare2.ed.ac.uk). Upon a pulse of light at CT16 or CT21 a statistical difference is observed between DD and pulsed *lak* larvae (****. p<0.0001 using a Mann-Whitney two-tailed test). In both phase delays (PD, pulse of light at CT16) and phase advance paradigms (PA. pulse of light at CT21), no statistical difference between *lak* and double larvae is observed using a Mann-Whitney two-tailed test. For each type of paradigm DD, PD and PA three independent experiments were pooled.

## Discussion

Experiments performed in mouse have established a role for melanopsin expressing cells of the eye in mediating light input to the circadian rhythm in mammals. In addition to the eye, non-mammalian vertebrates exhibit several extraocular sites of photoreception, raising the question of the relative impact of those different inputs. Herein we show that circadian rhythms of locomotor activity are established and photoentrain in the absence of RGCs in zebrafish larvae. Furthermore, our results show that the absence of a functional eye affects masking, but in a circadian dependent manner. As zebrafish also possesses melanopsin expressing cells in their pineal gland (Sapède et al., 2020), we engineered an *opn4xa* mutant line to address the role of *opn4xa*-dependent photosensitivity in this structure. Our data suggests that *opn4xa* is neither involved in masking nor in the establishment/photoentrainment of circadian rhythms. While our analysis does not support a redundant role for RGCs and *opn4xa* photosensitivity during photoentrainment of circadian rhythms it reveals a subtle function of *opn4xa*-dependent photosensitivity, possibly in the pineal, in the control of period length of circadian rhythms in constant light.

### Subtle defects in *opn4xa*-/- mutants in LL

While no differences in period or amplitude of locomotor rhythms are observed for *opn4xa* mutant larvae in DD, subtle alterations are observed in LL. Abrogation of *opn4xa* activity reduces the increase of period length observed when larvae are placed in LL. A similar defect is observed in Opn4- /- mice placed in constant light condition (Panda et al., 2002; Ruby et al., 2002) suggesting this could be a conserved function of melanopsin. Interestingly, this phenotype is not observed in lak mutant suggesting that in zebrafish this melanopsin function might involve the pineal gland rather than the eye. While the effect of *opn4xa* mutation on period length in LL is subtle, such an effect is not at all observed in DD. We see two possible hypotheses. First, *opn4xa* homozygous larvae could sense less light than controls and since period shows a reverse correlation with the intensity of light exposure (Aschoff, 1960; Ben-Moshe Livne et al., 2016), it could result in a lower period. A second hypothesis is that the *opn4xa* -/- clock is less stable in LL compared to DD. Indeed, we observed a stronger dampening of the circadian rhythms in LL compared to DD especially when four cycles were filmed and despite a moderate light intensity (20 lux). Along the same line, the variation observed for the periods are higher in LL than in DD for both control and mutant larvae suggesting again less robustness in clock activity in constant light conditions.

While we have analysed the expression of clock genes our results did not yield a clear molecular explanation for the *opn4xa* -/- period phenotype in LL. Indeed *opn4xa* -/- larvae only show a modest increase in Bmal1a, cry1 and dec1 expression in LL (Figure 5). A loss of dec1 using morpholino antisense technology suggests a role for this gene in controlling circadian rhythms of locomotor activity induced by a light pulse but a role for this gene in period control in LL has not been addressed (Ben-Moshe et al., 2014). Along the same line a double mutant line for per2 and cry1a exhibit deficits in the generation of rhythms following a light pulse but the role of cry1a during period control in LL has not been addressed (Hirayama et al., 2019) In contrast, per2 regulates period length in constant light but its transcription is not affected in *opn4xa*-/- larvae ((Ruggiero et al., 2021; Wang et al., 2015); Figure 5G)); it is however possible that affecting the level of CRY1A could modify PER2 activity therefore leading to the observed phenotype. Further experiments will be required to address this question.

### RGCs, but not *opn4xa*, are involved in masking

Compared to their control siblings, we found *lak* mutant larvae to be less active during the light phases of LD cycles as well as when subjected to a pulse of light at CT16 but not at CT21. This reveals a role for RGCs in positive masking in zebrafish larvae as well as a circadian control of this masking activity. It is important to note that while we refer to this as an effect of ‘masking’ it is at this step impossible to distinguish whether the *lak* mutation impairs the arousing effect of visual stimuli or the arousing effect of light itself. Moreover, masking is not completely abolished in *lak* mutant larvae: in LD as well as during a light pulse at CT16, *lak* mutant larvae still show some masking, albeit at a reduced level and masking is not affected in these larvae at CT21. Could *opn4xa* photosensitivity from the pineal compensate for the lack of RGCs? *opn4xa* -/- larvae display no defect in masking of locomotor activity in LD or during a pulse of light suggesting that *opn4xa*-dependent photosensitivity is dispensable for this type of masking. In addition, lak;opn4xa double mutants show a similar activity to *lak*-/- larvae during a pulse of light at CT16 and CT21, suggesting that there is no redundancy between the eye and *opn4xa*+ cells in the pineal for masking control. Other photosensitive cells are thus involved in this process. Among these could be the classical photoreceptors of the pineal or deep brain photoreceptors, such as those involved in the locomotor response to a loss of illumination (Fernandes et al., 2012).

### A Differential Role for RGCs in controlling phase advance versus phase delays in the rhythms of locomotor activity?

*lak* mutant larvae show a reduced phase-advance but no defect in a phase delay paradigm which could indicate a specific role for the RGC in controlling photoentrainment in a phase-advance context. On the other hand, *lak* mutant larvae display less aggregated pigment granules than siblings (Figure S5) Could this impact the way light penetrates their body? While this remains a possibility, we do not think it is a very likely hypothesis since 1.even in *lak* mutant there are rather wide pigment-free areas over the fish brain and in particular over the pineal (Figure S5B) 2.light can still penetrate from the side of the body as pigments are found only on the dorsal and ventral most epidermis (Figure S5C). Finally, experiments aiming at targeting different RGC subpopulations will help consolidate the role of the eye in controlling phase advances as well as identifying downstream circuits.

### RGCs and *opn4xa* are largely dispensable for shifting circadian rhythms of locomotor activity in response to a pulse of white light

The present study shows that neither the eye nor *opn4xa* mediated photosensitivity in the pineal gland is absolutely required for the development of circadian rhythms and circadian photoentrainment. The absence of a strong requirement for the eye to control the circadian system in zebrafish is surprising given that *Astyanax mexicanus* blind cavefishes are arrhythmic in DD (Beale et al., 2013) while *Phreatichthys andruzzii* adult cavefishes, which also exhibit a complete eye degeneration, and are arrhythmic in LD when fed at random times (Cavallari et al.. 2011). In light of our data, we propose that apart from the eye, other photosensitive structures might be affected in these other fish species. This in turn brings the question of which structure(s) relay light information to control circadian rhythm in fishes and other non-mammalian animals? The pineal gland, with its classical photoreceptors and *opn4xa*+ projection neurons is an appealing candidate (Sapède and Cau. 2013). Strategies aiming at genetically killing this structure or impairing its activity will surely help unravelling its function. Studies describing the effect of surgical pinealectomy have been reported in a number of non-mammalian vertebrates. The phenotypes induced seem to depend strongly on the species. For instance, pinealectomy abolishes rhythms in the stinging catfish but not in the amur catfish or the lake chub. Interestingly, in species where rhythms are maintained upon pinealectomy a change in period can occur (see (Zhdanova and Reebs, 2005) for a review). A similar variety of phenotypes are induced upon pinealectomy in reptiles or birds. In addition to the pineal gland, reptiles have a parietal eye, a structure that is developmentally and spatially related to the pineal gland. Interestingly simultaneous removal of the eye, the pineal gland and the parietal eye in two species of lizards (P. Sicula and S. olivaceous), does not impair rhythms of locomotor activity while on the contrary these rhythms are lost if in addition to this triple ablation injection of dark ink between the skin and the skull is performed (Tosini et al.. 2001). Similarly, experiments in songbirds suggest the existence of additional photosensitive structures located in the brain that control photoentrainment (Menaker and Underwood. 1976). Altogether these results highlight the existence of other brain structures mediating light inputs on the circadian system. Interestingly, melanopsin expression has been described in other brain areas in the zebrafish larva: opn4a is expressed within the presumptive optic area, opn4b is found in the ventral forebrain and the thalamic region, and opn4.1 is detected in a specific domain located in the ventricular region at the junction between the caudal hindbrain and the anterior spinal cord (Fernandes et al.. 2012; Matos-Cruz et al.. 2011).

Larvae double mutant for *opn4*.*1* and *opn4xb* show a decreased locomotor activity during the day but no circadian phenotype (Dekens et al., 2022). As 42 opsin genes are predicted in the zebrafish genome, of which 20 are expressed in the adult brain (Davies et al., 2015), further characterization of their expression in the larval brain will be needed to define the best candidates for further study. Finally, the possibility remains that photoentrainment in zebrafish occurs as a result of direct photosensitivity of motoneurons or muscles themselves as all cells and organs have been shown to be directly photosensitive and light-entrainable in this species (see (Vatine et al., 2011) for a review).

Taken together, our results highlight profound differences in the establishment and photoentrainment of the circadian system between the diurnal zebrafish and other species such as mice and human. A crucial, yet open question is whether these divergences reflect the different phylogeny of these species or their different use of temporal niches. The photosensitive capabilities of the zebrafish in particular and of aquatic species in general (as judged by the number of opsins predicted in the genome) far exceed that observed in mammals. This could imply a greater level of complexity and robustness in circadian control in zebrafish independently of its temporal niche. However, the human brain also expresses opsins (OPN3 and OPN5; (Halford et al., 2001; Tarttelin et al., 2003)) suggesting the existence of deep brain photoreceptors in diurnal primates and the possibility that they participate in photoentrainment.

## Acknowledgements

We are indebted to C. Rampon for allowing us to use the Ethovision Software as well as Julie Batut, Ouria Dkhissy Benyahya and Frederique Gaits-Iacovoni for helpful discussions. We thank M. Halpern and K. Soanes for sharing probes. This work was supported by the Centre National de la Recherche Scientifique (CNRS); the Institut National de la Santé et de la Recherche Médicale (INSERM); Université de Toulouse III (UPS); Fondation pour la Recherche Médicale (FRM; DEQ20131029166); Fédération pour la Recherche sur le Cerveau (FRC); Association pour la Recherche sur le Cancer (ARC); Association Rétina France and the Ministère de la Recherche. We would like to thank Brice Ronsin, Stéphanie Bosch and the Toulouse RIO Imaging platform; Stéphane Relexans, Aurore Laire and Richard Brimicombe for taking care of the fish as well as Sophie Polès for technical help.

## MATERIAL AND METHODS

### Zebrafish lines and developmental conditions

All animals were handled in the CBI fish facility, which is certified by the French Ministry of Agriculture (approval number A3155510). The project was approved by the French Ministry of Teaching and Research (agreement number APAFIS#3653-2.016.011.512.005.922). Embryos were reared at 28 degrees in a 14h light/10h dark cycle with lights on at 9:00 and lights off at 23:00. The *lak* mutant line has been described previously (Kay et al., 2001), *lak* homozygous mutants were identified by their dark coloration. The protocol for genotyping *lak* individuals is available upon request.

### Generation of an *opn4xa* mutant allele

An *opn4xa* mutant allele was generated using the CRISPR/ Cas9 targeted genome editing. For this. a target site was designed in the second exon by manual screening for PAM sites. Transcription of the guide and coinjection of the guide mRNA with cas9 mRNA was performed as described in (Lekk et al., 2019). Screening of potential mutants was performed using T7 endonuclease (NEB) treatment of PCR products amplifying the second *opn4xa* exon (Fw : 5’ CACAACATAAACTGTAACTGCATCC 3’, Rev : 5’ GACACGGGTATGACACTCAGGAAGG 3’). PCR products from potential carriers were subsequently subcloned and sequenced. In this manner we identified several interesting carriers among which an individual transmitting an allele bearing 17 extra nucleotides in the second exon leading to a premature interruption of the coding sequence.

To genotype *opn4xa* individuals, we used a classical PCR protocol with the following oligos: 5’-GGACGCCTCCAAACTTC-3’ (Forward) and 5’-CGAACACCCACTCCTTGTAC-3’ Reverse). PCR products of different sizes were obtained (110bp for the wt allele and 127bp for the mutant allele) and resolved on a 4% agarose gel.

### Locomotor Activity Assays

Larvae zebrafish coming from heterozygous incrosses were raised on a 14:10 hr light:dark cycle at 28°C in Petri dishes with no more than 50 larvae per Petri dish in a water bath inside the fish facility. On the morning of their 5 ^th^day of development (9:15-10:30), individual larvae were placed in each well of a 96-well plate containing aquarium fish water and placed back in the water bath. Light intensity during the entrainment was 110 lux at 6500K. On the evening (16:00-20:00), the plate was put in an hermetic box in a dark room maintained at 27°C with a heater. This home-made box (Vignet et al., 2013) was continuously illuminated from below with two panels of infra-red lights as well as neutral white light (4000K, 20 lux at water surface)) controlled by a timer from 9:00 to 23:00. Larvae were then filmed at 30 frames per second, with a ceiling mounted infra-red camera connected to a computer on the following days (from the 6^th^ day of development to the 9^th^ or 10^th^ day of development) in controlled conditions of illumination. The temperature inside the box was monitored using an electronic programmable device (I Button. Maxim). After the experiment, larvae were either genotyped by PCR for *opn4xa* and/or lak and/or simply identified for the lak mutation using the dark coloration phenotype. In addition, larvae presenting developmental defects were discarded from the subsequent analysis. Experiments in which too many larvae presented development issues or where temperature issues were present were discarded. At least three experiments were made for each type of assay.

### In Situ Hybridization

In situ hybridization was performed as described previously (Cau et al., 2008). *opn4xa*, tcf7 and c-fos probes have been described previously (Matos-Cruz et al.,2011; Ellis et al.,2012; Sapède et al., 2020).

### Locomotor Activity Analysis

After the experiment, the distance travelled per minute was extracted for each larva using the Ethovision XT13.0 (Noldus. Wageningen. the Netherlands) with the following parameters: for Detection Settings: dynamic substraction ; subject color compared to background : Darker ; Dark : 7 to 210 ; Frame Weight : 2 ; for Track Smoothing Profiles :Minimal Distance Moved : 0.2mm - Direct (A>MDM) ; for Data Profiles : Results per time bin. Ignore last time bin if incomplete; for Analysis Profiles: Distance moved of the center-point. The obtained files were then analysed using the wakefish program (written in python by L.Sanchou) to extract an average activity of mm/min over 10min for each larva (‘DM10 files’). For each experiment, the same number of homozygous mutants and wild-type or control larvae were randomly selected. The Biodare software was used to calculate periods and phases for each larva (biodare2.ed.ac.uk). We choose to use the FFT-NLLS to calculate periods and phases on DM10 files after baseline detrending, as advised (Zielinski et al., 2014). The parameters used for period calculation were as follows: baseline detrending, expected periods from 18 to 30 hours. analysis method FFT-NLLS. The parameters used for phase calculation were as follows: baseline detrending. FFT-NLLS, phase by fit, absolute phase to window. Windows used to calculate the phase of locomotor activity “before the pulse” and “after the pulse” encompass time points from CT0 to CT15 (corresponding from 9am of the 1^st^ day in of the experiment to midnight between the 2^nd^ and 3^rd^ day of the experiment for “before the pulse” and from 9am of the 3^rt^ day of the experiment to midnight between the 4^th^ and 5^th^ day of the experiment). Locomotor activity levels were calculated from the DM10 files by calculating means of the average activity in mm/min over 10 min over a given period for each larva. Statistical analysis was done using Prism. Graphs were generated using R studio (ggplot2 and rethomics packages (Geissmann et al., 2019; Wickham, 2016)).

### Analysis of clock gene expression using RTqPCR

For each specific stage, pools of 15 larvae were collected and extracted with TRIzol. Reverse transcription and qPCR were performed using a standard protocol (Quillien et al., 2021) with the oligos detailed in Supplemental Table 7. All the experiments were performed in triplicates and the mean expression relative to beta actin was calculated. Three pools of larvae were used for each time point.

## Supplemental data

**Supplemental Figure 1:**
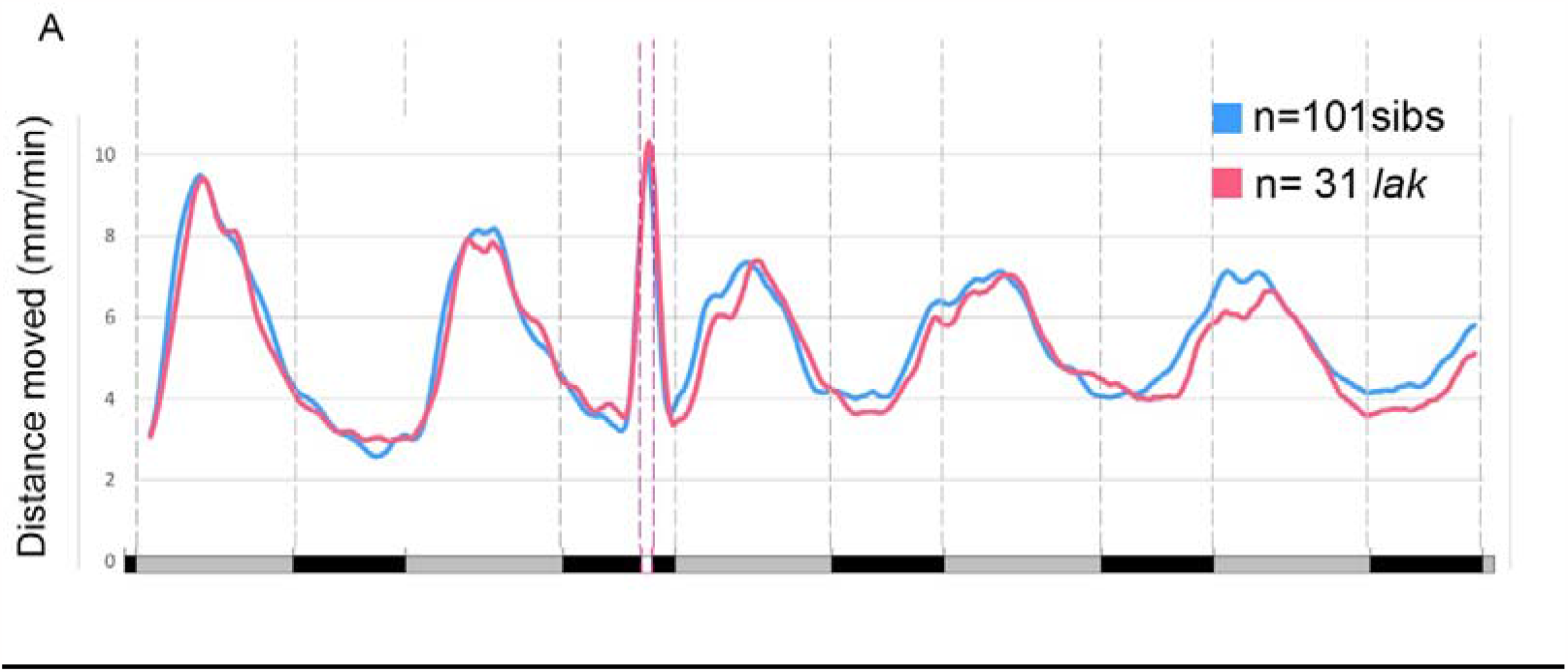
Figure S1 : (A) Average distance moved merged from PA experiments in 10 min bins. Mean ± SE. The original data is the same than in figure2 but here, only the larvae for which a phase could be extracted for the two first and the two last cycles were included in the average. In addition, two rounds of smoothing each using ten successive time points were applied as this made the difference in phase shift between the *lak* and the sib larvae more easy to visualize.

**Supplemental Figure 2:**
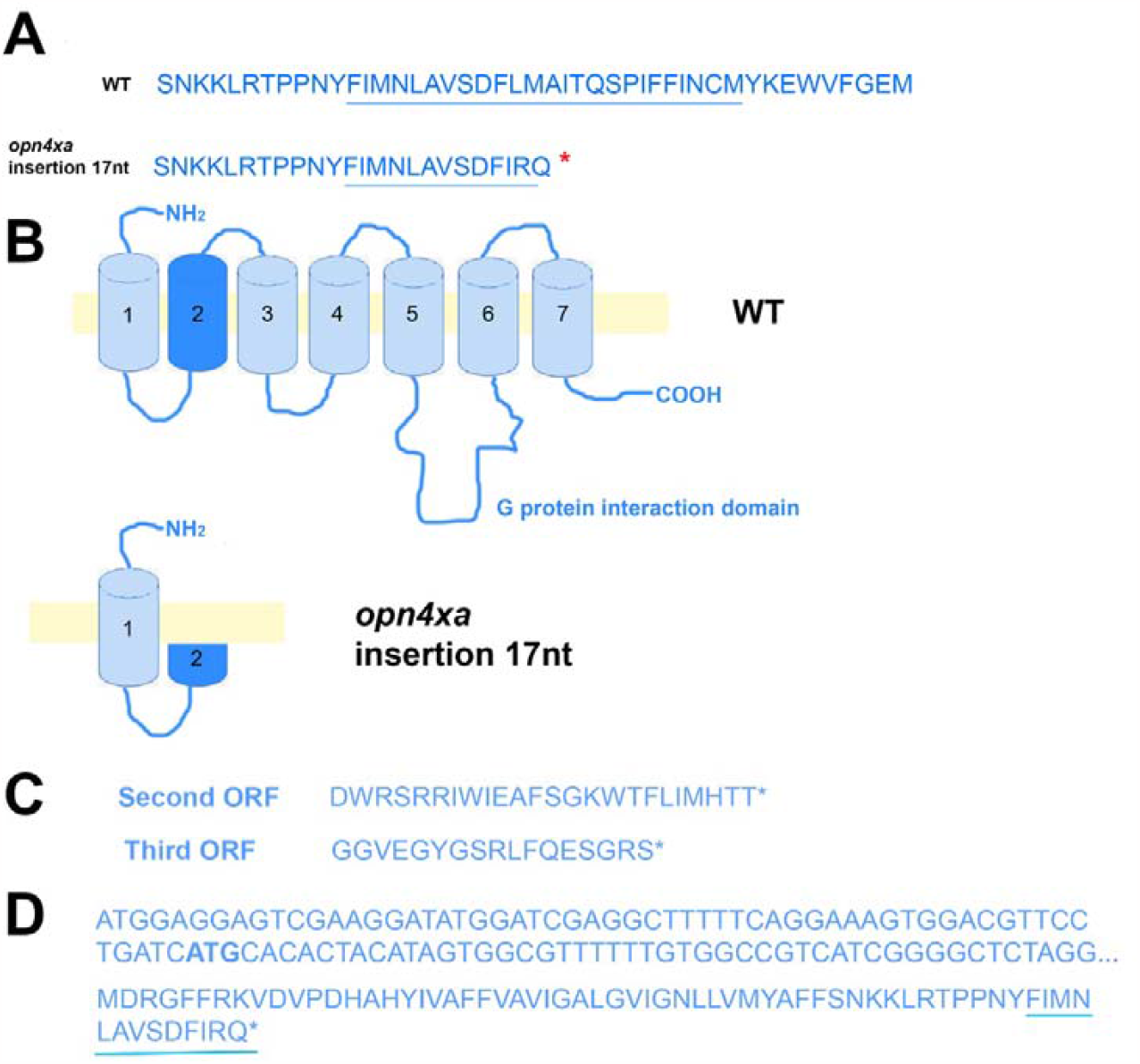
A)Prediction of the protein sequences produced by the wt and mutant exon 2. The part corresponding to the second transmembrane domain (Matos-Cruz et al., 2011) is underlined. The red asterisk indicates a premature stop codon. (B) Models of the WT and mutant predicted *OPN4XA* proteins.(C) Predicted proteins produced from alternative codon usage of the mutant cDNA. (D) cDNA sequence of the beginning of the *opn4xa* gene, a secondary ATG is indicated in bold. Below, the predicted protein produced from this alternative ATG in the mutant cDNA is indicated. In all cases described in C and D, the predicted mutant protein is truncated.

**Supplemental Figure 3:**
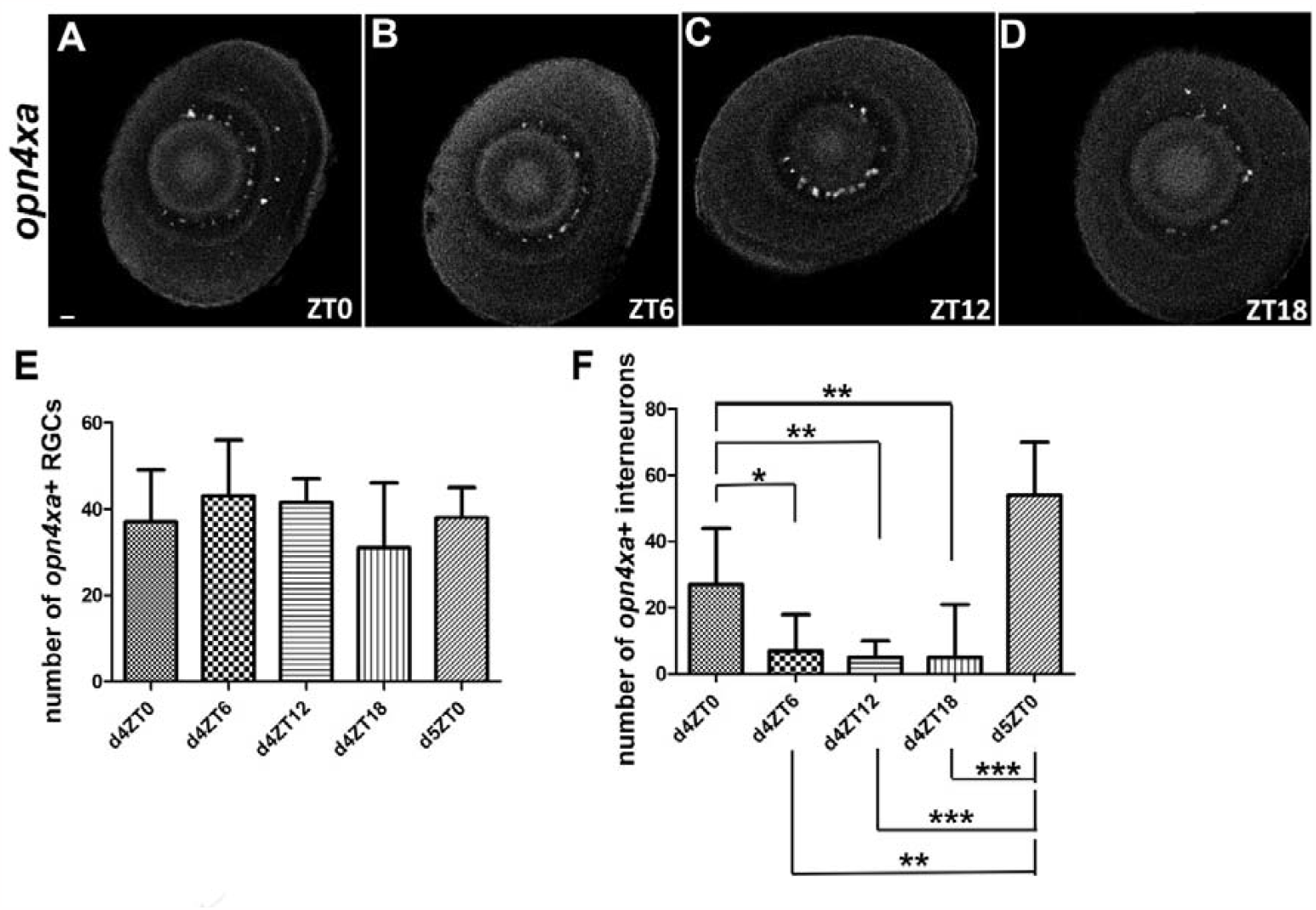
Figure S3: *opn4xa* is expressed in zebrafish RGCs and interneurons. **(A-D)** Expression of *opn4xa* at 4days at different ZT using fluorescent in situ hybridization. Lateral view of mounted eyes imaged under the confocal microscope. The ventral side is oriented towards the left upper corner. Based on position, the *opn4xa*+ cells from the interneuron layer are most likely horizontal and amacrine cells. **Scale bar: 10 µm.** **E)** Number of *opn4xa*+ cells in the RGC layer in 96-128 hpf zebrafish larvae. All data follows a Gaussian distribution. No statistical differences were observed between the different time points using a one-way ANOVA with Bonferroni post hoc test. **F)** Number of *opn4xa*+ cells in the interneuron layer in 96-128 hpf zebrafish larvae. The data at 4dZT0 does not follow a Gaussian distribution. * p<0.05. ** p<0.001. *** p<0.0005 using a Kruskal-Wallis test with Dunn’s post hoc comparison.

**Supplemental Figure 4:**
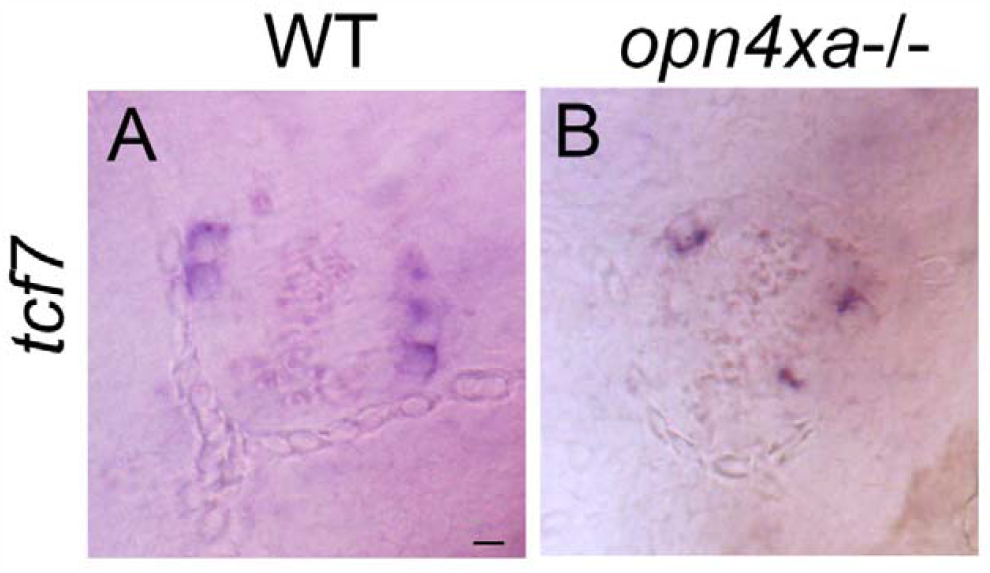
Figure S4: Expression of tcf7 in the pineal gland at 6 days using in situ hybridization in wt and *opn4xa*-/- larvae. Dorsal views are shown. Scale bar : 10 µm.

**Supplemental Figure 5:**
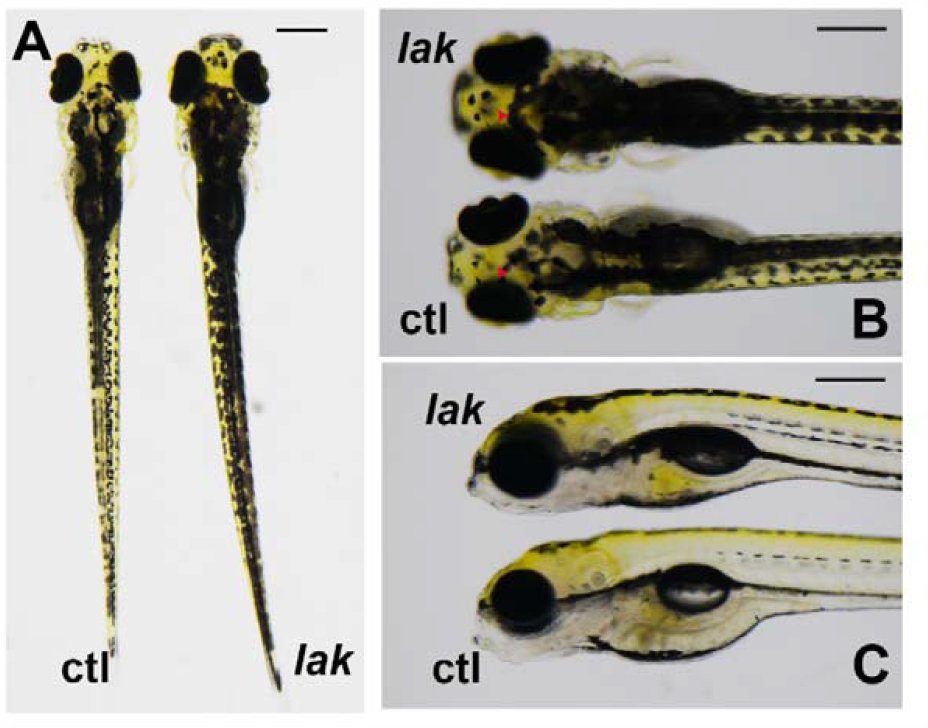
Figure S5 Live *lak* and sibling larvae at 5 dpf showing the differences in pigmentation. A-B) Dorsal views C) side views. The red arrow in B points to the position of the pineal gland which is not covered by pigments in *lak* and sib larvae. Scale bar: 0.5 mm.

## Legend for the following supplemental tables

### Legend for the supplemental tables S1-S6

Tables showing the average distance travelled (mm/min) over a 10 min window averaged during the day (D) or the night (N) periods. Mean ± S.D. D1 corresponds to the first day. The p value and statistical significance using a two-tailed Mann-Whitney test is indicated.

### Legend for the Supplemental table S7

List of primers for RTqPCR. The gene ID corresponds to the ZDB number.

**Supplemental table 1:** activity of *lakritz* -/- versus control larvae in LD.

| condition | ctrl (n=55) | *lakritz* (n=55) | p value |
| --- | --- | --- | --- |
| D1 | 13.95 ± 9.76 | 9.72 ± 8.73 | **\*\* 0.008** |
| N1 | 2.92 ± 2.28 | 2.85 ± 2.57 | n.s 0.42 |
| D2 | 16.90 ± 10.25 | 11.18 ± 7.40 | **\*\*\* 0.0005** |
| N2 | 4.04 ± 2.22 | 4.98 ± 4.22 | n.s 0.51 |
| D3 | 13.19 ± 8.56 | 10.88 ± 7.07 | n.s 0.13 |
| N3 | 3.56 ± 2.17 | 3.95 ± 2.94 | n.s 0.57 |

**Supplemental table 2:** activity of *lakritz* -/- versus control larvae in DD.

| condition | ctrl (n=84) | *lakritz* (n=84) | p value |
| --- | --- | --- | --- |
| D1 | 8.9 ± 8.7 | 8.73 ± 7.08 | n.s 0.43 |
| N1 | 2.78 ± 3.4 | 2.45 ± 1.67 | n.s 0.98 |
| D2 | 6.20 ± 4.18 | 6.39 ± 5.77 | n.s 0.44 |
| N2 | 2.64 ± 1.60 | 2.80 ± 5.64 | n.s 0.92 |
| D3 | 6.76 ± 7.25 | 6.86 ± 5.64 | n.s 0.95 |
| N3 | 3.13 ± 1.95 | 3.27 ± 2.16 | n.s 0.87 |
| D4 | 8.68 ± 8.41 | 8.52 ± 6.89 | n.s 0.55 |
| N4 | 3.42 ± 2.20 | 3.43 ± 2.18 | n.s 0.92 |

**Supplemental table 3:** activity of l*akritz* -/- versus control larvae in LL.

| condition | ctrl (n=72) | *lakritz* (n=72) | p value |
| --- | --- | --- | --- |
| D1 | 21.31 ± 13.33 | 19.49 ± 12.39 | n.s 0.44 |
| N1 | 7.38 ± 4.59 | 6.61 ± 4.83 | n.s 0.27 |
| D2 | 22.99 ± 14.01 | 22.88 ± 19.03 | n.s 0.27 |
| N2 | 8.03 ± 4.35 | 9.28 ± 6.59 | n.s 0.35 |
| D3 | 17.71 ± 8.51 | 17.43 ± 14.59 | n.s 0.096 |
| N3 | 8.11 ± 3.63 | 8.54 ± 5.38 | n.s 0.85 |

**Supplemental table 4:** activity of *opn4xa* -/- versus control larvae in LD.

| condition | wt (n=48) | *opn4xa* *-/-* (n=48) | p value |
| --- | --- | --- | --- |
| D1 | 22.16 ± 10.02 | 22.42 ± 13.12 | n.s 0.73 |
| N1 | 9.41 ± 6.45 | 7.94 ± 4.99 | n.s 0.30 |
| D2 | 20.46 ± 7.96 | 19.49 ± 8.25 | n.s 0.50 |
| N2 | 7.27 ± 4.86 | 6.32 ± 4.53 | n.s 0.27 |
| D3 | 14.35 ± 5.19 | 12.52 ± 6.12 | n.s 0.07 |
| N3 | 5.59 ± 3.41 | 4.19 ± 3.09 | **\* 0.01** |

**Supplemental table 5:**
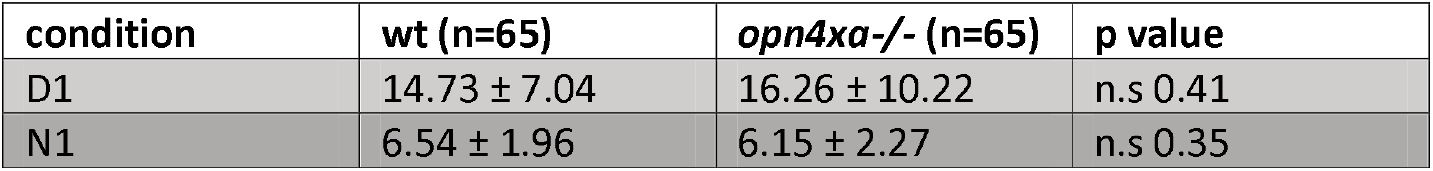
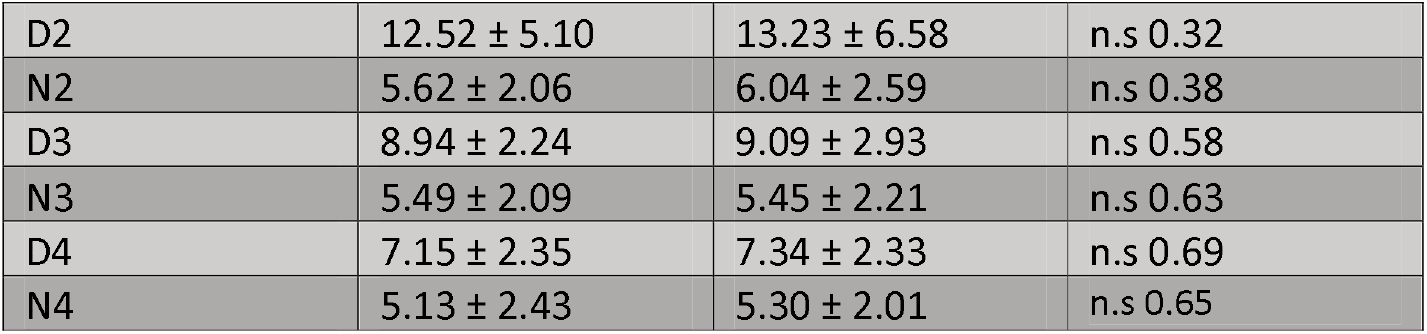
activity of *opn4xa* -/- versus control larvae in DD.

| condition | wt (n=65) | *opn4xa* *-/-* (n=65) | p value |
| --- | --- | --- | --- |
| D1 | 14.73 ± 7.04 | 16.26 ± 10.22 | n.s 0.41 |
| N1 | 6.54 ± 1.96 | 6.15 ± 2.27 | n.s 0.35 |
| D2 | 12.52 ± 5.10 | 13.23 ± 6.58 | n.s 0.32 |
| N2 | 5.62 ± 2.06 | 6.04 ± 2.59 | n.s 0.38 |
| D3 | 8.94 ± 2.24 | 9.09 ± 2.93 | n.s 0.58 |
| N3 | 5.49 ± 2.09 | 5.45 ± 2.21 | n.s 0.63 |
| D4 | 7.15 ± 2.35 | 7.34 ± 2.33 | n.s 0.69 |
| N4 | 5.13 ± 2.43 | 5.30 ± 2.01 | n.s 0.65 |

**Supplemental table 6:** activity of *opn4xa* -/- versus control larvae in LL.

| condition | wt (n=66) | *opn4xa* -/- (n=66) | p value |
| --- | --- | --- | --- |
| D1 | 16.56 ± 9.57 | 20.15 ± 12.25 | n.s 0.07 |
| N1 | 3.84 ± 2.99 | 6.22 ± 5.60 | * **0.02** |
| D2 | 16.67 ± 7.79 | 19.22 ± 11.49 | n.s 0.26 |
| N2 | 4.90 ± 3.30 | 5.92 ± 4.48 | n.s 0.36 |
| D3 | 14.27 ± 7.16 | 13.94 ± 7.26 | n.s 0.82 |
| N3 | 5.92 ± 4.09 | 5.63 ± 3.75 | n.s 0.70 |

**Supplemental table 7:**
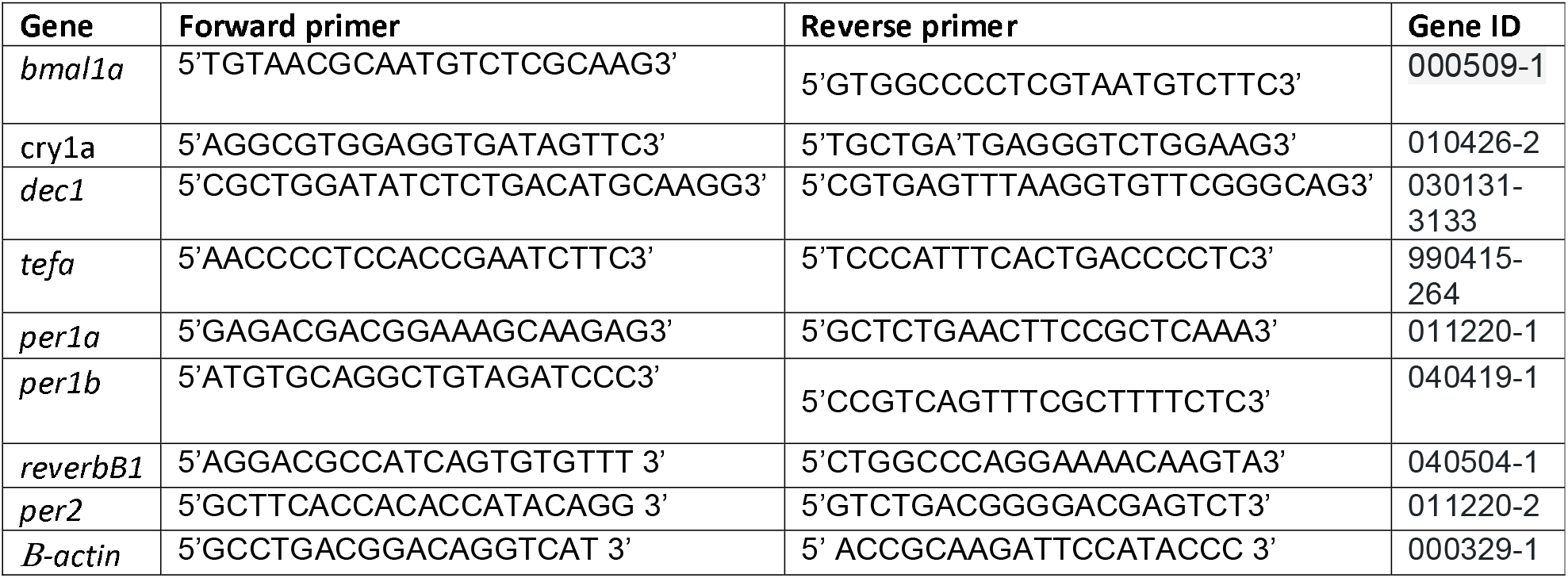
*qRTPCR* primer sequences.

## REFERENCES

Altimus, C. M., Güler, A. D., Villa, K. L., McNeill, D. S., Legates, T. A. and Hattar, S. (2008). Rods-cones and melanopsin detect light and dark to modulate sleep independent of image formation. Proc. Natl. Acad. Sci. U. S. A. 105, 19998–20003.

Aschoff, J. (1960). Exogenous and endogenous components in circadian rhythms. Cold Spring Harb. Symp. Quant. Biol. 25, 11–28.

Baver, S. B., Pickard, G. E., Sollars, P. J. and Pickard, G. E. (2008). Two types of melanopsin retinal ganglion cell differentially innervate the hypothalamic suprachiasmatic nucleus and the olivary pretectal nucleus. Eur. J. Neurosci. 27, 1763–1770.

Beale, A., Guibal, C., Tamai, T. K., Klotz, L., Cowen, S., Peyric, E., Reynoso, V. H., Yamamoto, Y. and Whitmore, D. (2013). Circadian rhythms in Mexican blind cavefish Astyanax mexicanus in the lab and in the field. Nat. Commun. 4, 1–10.

Belenky, M. A., Smeraski, C. A., Provencio, I., Sollars, P. J. and Pickard, G. E. (2003). Melanopsin retinal ganglion cells receive bipolar and amacrine cell synapses. J. Comp. Neurol. 460, 380–393.

Ben-Moshe, Z., Alon, S., Mracek, P., Faigenbloom, L., Tovin, A., Vatine, G. D., Eisenberg, E., Foulkes, N. S. and Gothilf, Y. (2014). The light-induced transcriptome of the zebrafish pineal gland reveals complex regulation of the circadian clockwork by light. Nucleic Acids Res. 42, 3750–3767.

Ben-Moshe Livne, Z., Alon, S., Vallone, D., Bayleyen, Y., Tovin, A., Shainer, I., Nisembaum, L. G., Aviram, I., Smadja-Storz, S., Fuentes, M., et al. (2016). Genetically Blocking the Zebrafish Pineal Clock Affects Circadian Behavior. PLoS Genet. 12, e1006445.

Bhadra, B., N, T., P, D. and M, P. B. (2017). Evolution of circadian rhythms: from bacteria to human. Sleep Med.

Brzezinski, J. A., Brown, N. L., Tanikawa, A., Bush, R. A., Sieving, P. A., Vitaterna, M. H., Takahashi, J. S. and Glaser, T. (2005). Loss of Circadian Photoentrainment and Abnormal Retinal Electrophysiology in Math5 Mutant Mice. Invest. Ophthalmol. Vis. Sci. 46, 2540–2551.

Calligaro, H., Coutanson, C., Najjar, R. P., Mazzaro, N., Cooper, H. M., Haddjeri, N., Felder-Schmittbuhl, M.-P. and Dkhissi-Benyahya, O. (2019). Rods contribute to the light-induced phase shift of the retinal clock in mammals. PLoS Biol. 17, e2006211.

Carr, A.-J. F. and Whitmore, D. (2005). Imaging of single light-responsive clock cells reveals fluctuating free-running periods. Nat. Cell Biol. 7, 319–321.

Cau, E., Quillien, A. and Blader, P. (2008). Notch resolves mixed neural identities in the zebrafish epiphysis. Dev. Camb. Engl. 135, 2391–2401.

Cavallari, N., Frigato, E., Vallone, D., Fröhlich, N., Lopez-Olmeda, J. F., Foà, A., Berti, R., Sánchez-Vázquez, F. J., Bertolucci, C. and Foulkes, N. S. (2011). A Blind Circadian Clock in Cavefish Reveals that Opsins Mediate Peripheral Clock Photoreception. PLoS Biol. 9,.

Chew, K. S., Renna, J. M., McNeill, D. S., Fernandez, D. C., Keenan, W. T., Thomsen, M. B., Ecker, J. L., Loevinsohn, G. S., VanDunk, C., Vicarel, D. C., et al. (2017). A subset of ipRGCs regulates both maturation of the circadian clock and segregation of retinogeniculate projections in mice. eLife 6, e22861.

Covello, G., Rossello, F. J., Filosi, M., Gajardo, F., Duchemin, A., Tremonti, B. F., Eichenlaub, M., Polo, J. M., Powell, D., Ngai, J., et al. (2020). Transcriptome analysis of the zebrafish atoh7−/− Mutant, lakritz, highlights Atoh7-dependent genetic networks with potential implications for human eye diseases. FASEB BioAdvances 2, 434–448.

Davies, W. I. L., Tamai, T. K., Zheng, L., Fu, J. K., Rihel, J., Foster, R. G., Whitmore, D. and Hankins, M. W. (2015). An extended family of novel vertebrate photopigments is widely expressed and displays a diversity of function. Genome Res. 25, 1666–1679.

Dekens, M. P. S., Fontinha, B. M., Gallach, M., Pflügler, S. and Tessmar-Raible, K. (2022). Melanopsin elevates locomotor activity during the wake state of the diurnal zebrafish. EMBO Rep. 23, e51528.

Dkhissi-Benyahya, O., Gronfier, C., De Vanssay, W., Flamant, F. and Cooper, H. M. (2007). Modeling the role of mid-wavelength cones in circadian responses to light. Neuron 53, 677–687.

Dollet, A., Albrecht, U., Cooper, H. M. and Dkhissi-Benyahya, O. (2010). Cones Are Required for Normal Temporal Responses to Light of Phase Shifts and Clock Gene Expression. Chronobiol. Int. 27, 768–781.

Fernandes, A. M., Fero, K., Arrenberg, A. B., Bergeron, S. A., Driever, W. and Burgess, H. A. (2012). Deep brain photoreceptors control light seeking behavior in zebrafish larvae. Curr. Biol. CB 22, 2042–2047.

Fernandez, D. C., Chang, Y.-T., Hattar, S. and Chen, S.-K. (2016). Architecture of retinal projections to the central circadian pacemaker. Proc. Natl. Acad. Sci. 113, 6047–6052.

Freedman, M. S., Lucas, R. J., Soni, B., Schantz, M. von, Muñoz, M., David-Gray, Z. and Foster, R. (1999). Regulation of Mammalian Circadian Behavior by Non-rod, Non-cone, Ocular Photoreceptors. Science.

Geissmann, Q., Rodriguez, L. G., Beckwith, E. J. and Gilestro, G. F. (2019). Rethomics: An R framework to analyse high-throughput behavioural data. PLOS ONE 14, e0209331.

Gompf, H. S., Fuller, P. M., Hattar, S., Saper, C. B. and Lu, J. (2015). Impaired circadian photosensitivity in mice lacking glutamate transmission from retinal melanopsin cells. J. Biol. Rhythms 30, 35–41.

Güler, A. D., Ecker, J. L., Lall, G. S., Haq, S., Altimus, C. M., Liao, H.-W., Barnard, A. R., Cahill, H., Badea, T. C., Zhao, H., et al. (2008). Melanopsin cells are the principal conduits for rod-cone input to non-image-forming vision. Nature 453, 102–105.

Halford, S., Freedman, Melanie S., Bellingham, J., Inglis, S. L., Poopalasundaram, S., Soni, B. G., Foster, R. G. and Hunt, D. M. (2001). Characterization of a Novel Human Opsin Gene with Wide Tissue Expression and Identification of Embedded and Flanking Genes on Chromosome 1q43. Genomics 72, 203–208.

Hartley, S., Dauvilliers, Y. and Quera-Salva, M.-A. (2018). Circadian Rhythm Disturbances in the Blind. Curr. Neurol. Neurosci. Rep. 18, 65.

Hirayama, J., Alifu, Y., Hamabe, R., Yamaguchi, S., Tomita, J., Maruyama, Y., Asaoka, Y., Nakahama, K.-I., Tamaru, T., Takamatsu, K., et al. (2019). The clock components Period2, Cryptochrome1a, and Cryptochrome2a function in establishing light-dependent behavioral rhythms and/or total activity levels in zebrafish. Sci. Rep. 9, 196.

Karnas, D., Hicks, D., Mordel, J., Pévet, P. and Meissl, H. (2013). Intrinsic Photosensitive Retinal Ganglion Cells in the Diurnal Rodent, Arvicanthis ansorgei. PLOS ONE 8, e73343.

Kay, J. N., Finger-Baier, K. C., Roeser, T., Staub, W. and Baier, H. (2001). Retinal Ganglion Cell Genesis Requires lakritz, a Zebrafish atonal Homolog. Neuron 30, 725–736.

Kofuji, P., Mure, L. S., Massman, L. J., Purrier, N., Panda, S. and Engeland, W. C. (2016). Intrinsically Photosensitive Retinal Ganglion Cells (ipRGCs) Are Necessary for Light Entrainment of Peripheral Clocks. PLOS ONE 11, e0168651.

Kölsch, Y., Hahn, J., Sappington, A., Stemmer, M., Fernandes, A. M., Helmbrecht, T. O., Lele, S., Butrus, S., Laurell, E., Arnold-Ammer, I., et al. (2021). Molecular classification of zebrafish retinal ganglion cells links genes to cell types to behavior. Neuron 109, 645–662.e9.

Lekk, I., Duboc, V., Faro, A., Nicolaou, S., Blader, P. and Wilson, S. W. (2019). Sox1a mediates the ability of the parapineal to impart habenular left-right asymmetry. eLife 8, e47376.

Li, J. Y. and Schmidt, T. M. (2018). Divergent projection patterns of M1 ipRGC subtypes. J. Comp. Neurol. 526, 2010–2018.

Lockley, S. W., Arendt, J. and Skene, D. J. (2007). Visual impairment and circadian rhythm disorders. Dialogues Clin. Neurosci. 9, 301–314.

Masai, I., Stemple, D. L., Okamoto, H. and Wilson, S. W. (2000). Midline Signals Regulate Retinal Neurogenesis in Zebrafish. Neuron 27, 251–263.

Matos-Cruz, V., Blasic, J., Nickle, B., Robinson, P. R., Hattar, S. and Halpern, M. E. (2011). Unexpected diversity and photoperiod dependence of the zebrafish melanopsin system. PloS One 6, e25111.

Menaker, M. and Underwood, H. (1976). Extraretinal Photoreception in Birds. Photochem. Photobiol. 23, 299–306.

Mure, L. S. (2021). Intrinsically Photosensitive Retinal Ganglion Cells of the Human Retina. Front. Neurol. 12, 636330.

Neuhauss, S. C. F., Biehlmaier, O., Seeliger, M. W., Das, T., Kohler, K., Harris, W. A. and Baier, H. (1999). Genetic Disorders of Vision Revealed by a Behavioral Screen of 400 Essential Loci in Zebrafish. J. Neurosci. 19, 8603–8615.

Panda, S., Sato, T. K., Castrucci, A. M., Rollag, M. D., DeGrip, W. J., Hogenesch, J. B., Provencio, I. and Kay, S. A. (2002). Melanopsin (Opn4) requirement for normal light-induced circadian phase shifting. Science 298, 2213–2216.

Perez-Leon, J. A., Warren, E. J., Allen, C. N., Robinson, D. W. and Brown, R. L. (2006). Synaptic inputs to retinal ganglion cells that set the circadian clock. Eur. J. Neurosci. 24, 1117–1123.

Quillien, A., Gilbert, G., Boulet, M., Ethuin, S., Waltzer, L. and Vandel, L. (2021). Prmt5 promotes vascular morphogenesis independently of its methyltransferase activity. PLoS Genet. 17,.

Ruby, N. F., Brennan, T. J., Xie, X., Cao, V., Franken, P., Heller, H. C. and O’Hara, B. F. (2002). Role of melanopsin in circadian responses to light. Science 298, 2211–2213.

Ruggiero, G., Ben-Moshe Livne, Z., Wexler, Y., Geyer, N., Vallone, D., Gothilf, Y. and Foulkes, N. S. (2021). Period 2: A Regulator of Multiple Tissue-Specific Circadian Functions. Front. Mol. Neurosci. 14, 718387.

Rupp, A. C., Ren, M., Altimus, C. M., Fernandez, D. C., Richardson, M., Turek, F., Hattar, S. and Schmidt, T. M. (2019). Distinct ipRGC subpopulations mediate light’s acute and circadian effects on body temperature and sleep. eLife 8, e44358.

Sack, R. L., Lewy, A. J., Blood, M. L., Keith, L. D. and Nakagawa, H. (1992). Circadian rhythm abnormalities in totally blind people: incidence and clinical significance. J. Clin. Endocrinol. Metab. 75, 127–134.

Sapède, D. and Cau, E. (2013). The pineal gland from development to function. Curr. Top. Dev. Biol. 106, 171–215.

Sapède, D., Chaigne, C., Blader, P. and Cau, E. (2020). Functional heterogeneity in the pineal projection neurons of zebrafish. Mol. Cell. Neurosci. 103, 103468.

Tamai, T. K., Young, L. C. and Whitmore, D. (2007). Light signaling to the zebrafish circadian clock by Cryptochrome 1a. Proc. Natl. Acad. Sci. U. S. A. 104, 14712–14717.

Tarttelin, E. E., Bellingham, J., Hankins, M. W., Foster, R. G. and Lucas, R. J. (2003). Neuropsin (Opn5): a novel opsin identified in mammalian neural tissue 1. FEBS Lett. 554, 410–416.

Tosini, G., Bertolucci, C. and Foà, A. (2001). The circadian system of reptiles: a multioscillatory and multiphotoreceptive system. Physiol. Behav. 72, 461–471.

Vallone, D., Gondi, S. B., Whitmore, D. and Foulkes, N. S. (2004). E-box function in a period gene repressed by light. Proc. Natl. Acad. Sci. U. S. A. 101, 4106–4111.

Vatine, G., Vallone, D., Gothilf, Y. and Foulkes, N. S. (2011). It’s time to swim! Zebrafish and the circadian clock. FEBS Lett. 585, 1485–1494.

Wang, M., Zhong, Z., Zhong, Y., Zhang, W. and Wang, H. (2015). The Zebrafish Period2 Protein Positively Regulates the Circadian Clock through Mediation of Retinoic Acid Receptor (RAR)-related Orphan Receptor α (Rorα). J. Biol. Chem. 290, 4367–4382.

Wee, R., Castrucci, A. M., Provencio, I., Gan, L. and Van Gelder, R. N. (2002). Loss of Photic Entrainment and Altered Free-Running Circadian Rhythms in math5−/− Mice. J. Neurosci. 22, 10427–10433.

Whitmore, D., Foulkes, N. S. and Sassone-Corsi, P. (2000). Light acts directly on organs and cells in culture to set the vertebrate circadian clock. Nature 404, 87–91.

Wickham, H. (2016). ggplot2: Elegant Graphics for Data Analysis. Springer-Verl. N. Y. ISBN 978-3-319-24277-4.

Wong, K. Y., Dunn, F. A., Graham, D. M. and Berson, D. M. (2007). Synaptic influences on rat ganglion-cell photoreceptors. J. Physiol. 582, 279–296.

Zhdanova, I. V. and Reebs, S. G. (2005). Circadian Rhythms in Fish. Fish Physiol. Acad. Press 197–238.

Zielinski, T., Moore, A. M., Troup, E., Halliday, K. J. and Millar, A. J. (2014). Strengths and Limitations of Period Estimation Methods for Circadian Data. PLOS ONE 9, e96462.

